# Uncovering the “ZIP code” for bZIP dimers reveals novel motifs, regulatory rules and one billion years of *cis*-element evolution

**DOI:** 10.1101/2022.04.17.488518

**Authors:** Miaomiao Li, Wanru Lin, Will Hinckley, Tao Yao, Wellington Muchero, Jin-Gui Chen, S. Carol Huang

## Abstract

Many eukaryotic transcription factors (TF) form homodimer or heterodimer complexes to regulate gene expression. For example, dimerization properties of the basic leucine zipper (bZIP) family play a critical role in regulating the unique biological functions in all eukaryotes. However, the molecular mechanism underlying the binding sequence and functional specificity of homo- *versus* heterodimers remains elusive. To fill this gap, we developed a double DNA Affinity Purification sequencing (dDAP-seq) technique that maps heterodimer DNA binding sites in an endogenous genome context. Our genome-wide binding profiles of twenty pairs of C/S1 bZIP heterodimers and S1 homodimers in *Arabidopsis* revealed that heterodimerization significantly expands the DNA binding preferences of bZIP TFs. Analysis of the heterodimer target genes in stress response and development suggest heterodimerization gives rise to regulatory responses that are distinct from the homodimers. In addition to the classic ACGT elements recognized by plant bZIPs, we found that the C/S1 heterodimers bound to motifs that might share an origin with the GCN4 *cis*-elements in yeast that diverged from plants more than one billion years ago. Importantly, heterodimer binding specificities can be distinguished by their relative preference for ACGT motifs *versus* GCN4-related motifs. More broadly, our study demonstrates the potential of dDAP-seq in deciphering the DNA binding specificities of interacting TFs that are key for combinatorial gene regulation.

## Introduction

Sequence-specific transcription factors (TF) regulate the expression of their target genes by recognizing short DNA sequences known as transcription factor binding sites (TFBS). The specificity of TF-DNA interactions can also be affected by genome and epigenome context or protein cofactors^1–6^. In particular, most TFs work within a complex or interact with other proteins to regulate gene expression *in vivo*^7^. Interactions of TFs with other TFs could alter, enhance or repress DNA binding activity depending on their cooperative, synergistic or competitive relationships^7–9^. Importantly, for many TFs across diverse structural families, TF interactions modify the recognition motifs of each individual TFs, including alternate motif spacing and orientation^10^. Therefore, elucidating how the DNA binding specificity of a TF is altered by interaction with another TF is key to accurate understanding of combinatorial gene regulation and TF functions.

Homo- and heterodimerization is an important feature of DNA recognition and regulatory function for many TFs^1,11,12^. A well-known example in plants, the Auxin Response Factor (ARF) family of TFs, requires dimerization to bind DNA efficiently. Structural analysis supports a model where ARF dimers interact with DNA in multiple configurations, and the DNA binding affinity of a dimer can be much higher than that of each of the two monomers^3^. In another case, the basic helix-loop-helix (bHLH) TFs bind DNA as homodimers, while heterodimerization between HLH and bHLH inhibits its DNA binding activity^13–15^. Therefore, heterodimerization not only increases the combinatorial complexity for a limited number of TFs, but also enhances their functional specificity^16,17^.

The basic leucine zipper (bZIP) proteins are dimerizing TFs found in all eukaryotes^16^. Dimerization of the DNA-binding domain occurs via the leucine zipper domain that positions the “basic” region of each monomer in contact with DNA^18^. The expansion of the bZIP family by genome duplication appears to have driven a strong tendency for homo- and heterodimerization between members, forming a bZIP dimer-mediated regulatory network^16,19^. The dimerization properties of this family play a significant role in regulating the overlapping and unique biological functions of the family members in both plants and animals. For instance, the human TF AP-1 is formed by a mixture of homo- and heterodimers of JUN and FOS proteins where the dimer composition defines the pattern of target gene expression^20^. The human bZIP ATF3 acts as a repressor as a homodimer but becomes an activator when it heterodimerizes with JUN^21^. A systematic survey of DNA binding specificities of human bZIP heterodimers revealed that many heterodimers targeted new types of DNA binding motifs that were not bound by either of the interacting partners^11^. In plants, studies started in the 1990s explored the DNA binding activities of different bZIPs using a gel-shift approach which provided a foundation for subsequent investigations of DNA sequence recognition by plant bZIPs^22–24^. Among the key findings were that plant bZIPs preferred *cis-*elements containing the ACGT sequence, such as the G-box (CACGTG), C-box (CACGTC), and A-box (TACGTA), and nucleotides flanking these ACGT elements played important roles in determining specificity. In addition to the ACGT elements, a handful of other *cis*-acting elements, such as the ACT element (ACTCAT/ATGAGT), have also been identified as important binding sites for heterodimers formed by plant Group C and S1 bZIPs^25^.

In the model plant *Arabidopsis thaliana*, 78 bZIP proteins are classified into thirteen phylogenetic groups^26^. Heterodimerization has been widely observed among the paralogous groups C and S1^27,28^. Genetic studies revealed that combinatorial regulation by these two groups is essential to maintain plant growth and development, as well as response to biotic and abiotic stresses, such as in low energy environments^29–32^. For example, during nutrient starvation response, bZIP53 (Group S1) and bZIP10 (Group C) interact to strongly enhance the expression of *proline dehydrogenase* (*ProDH*), which catalyzes the breakdown of proline as part of amino acid recycling to support energy demand^29,33^. This activation is mediated by binding of the bZIP53 and bZIP10 heterodimer to the *ProDH* promoter at an ACT element that resembles a preferred binding sequence of GCN4, a well characterized bZIP from unicellular yeast that diverged from *Arabidopsis* one billion years ago^34–36^. *Asparagine synthetase 1* (*ASN1*), which encodes a glutamine-dependent central enzyme in asparagine synthesis^37^, is also regulated by the C/S1 heterodimer via C-box, G-box and ACT motifs present in its promoter region^29,30,37,38^. These findings suggest that bZIP C/S1 heterodimerization gives rise to important functions and regulatory mechanisms that are distinct from the homodimers.

Despite the importance of heterodimerization to the functions of the C/S1 bZIPs, the molecular mechanism underlying the functional diversity and specificity of different homo- and heterodimers remains elusive. The pairs interact promiscuously in several assays^27,28^ and have a high degree of co-expression across tissue types and conditions^39^, so neither protein-protein interaction nor co-expression patterns are sufficient to explain the unique functions for the heterodimers. Here we extended our previously published method DAP-seq (DNA Affinity Purification sequencing)^40,41^, which identifies genome-wide TFBS for single TFs, to allow mapping of binding sites on endogenous genomic DNA for interacting TFs. Using this method, we generated binding site maps for twenty pairs of the C/S1 homo- and heterodimers. We found that a substantial number of binding sites for the heterodimers were not shared with the homodimers, and that these unique binding events were associated with novel transcriptional responses and predicted biological processes. Importantly, the heterodimer- and homodimer-specific binding events could be distinguished by the presence of the classic ACGT elements *vs*. the GCN4-related elements. Compared to experiments that interrogated binding of interacting TFs on synthetic oligonucleotides that contained limited genomic context^10,11^, our binding site maps captured the full complexity and sequence representation of real genomic DNA. The rules that we uncovered for combinatorial TF binding would provide valuable insights for understanding combinatorial gene regulation.

## Results

### Genome-wide binding profiles of bZIP C/S1 homo- and heterodimers by dDAP-seq

The bZIP family in *Arabidopsis* consists of thirteen phylogenetic groups and 78 members. Functional characterization is challenging due to the high degree of redundancy and synergistic interactions among family members^26,42^. We sought to deconvolute some of this complexity by studying the genome-wide binding profiles of the homodimers and heterodimers formed by members in the groups C and S1. We hypothesized that the functional specificities could be mediated by variation of DNA binding specificities between the pairs, which had not been investigated systematically due to technical limitations. Previously we published the DAP-seq method that used *in vitro* expressed TFs to interrogate DNA libraries constructed from naked genomic DNA^40,41^. Compared to methods that assay protein binding on synthetic oligonucleotides, such as systematic evolution of ligands by exponential enrichment (SELEX) and protein binding microarray (PBM)^10,43^, DAP-seq binding events occur in the context of endogenous genome sequence and DNA chemical modification. In contrast to *in vivo* methods such as chromatin immunoprecipitation followed by sequencing (ChIP-seq), which finds but does not distinguish direct and indirect binding events^44,45^, DAP-seq reports binding sites from direct interaction between the expressed TF and genomic DNA. We first applied DAP-seq to Group S1 members (bZIP1, bZIP2, bZIP11, bZIP44, bZIP53; Fig.1a) and in agreement with their previously described capacity to bind DNA as homodimers^28,42^, we successfully identified 5,200 to 22,000 binding events genome-wide for each of the members (Fig. 1b). In contrast, no binding site enrichment was found for any of the four Group C members (bZIP9, bZIP10, bZIP25, bZIP63) when tested alone using the same DAP-seq strategy despite the presence of a highly conserved bZIP DNA-binding domain (Fig. 1b). To ensure our inability to detect DNA-binding of Group C members was not caused by poor protein expression or protein misfolding, we performed *in vitro* pull-down assays using similar conditions to DAP-seq and detected protein-protein interactions between Group C and Group S1 bZIPs that were consistent with published results from *in vivo* assays^27,28^ (Supplementary Fig.1a-d). These data together suggested that Group C bZIPs, while unable to bind DNA on their own, might form functional DNA-binding partners with Group S1 bZIPs, and motivated us to develop a new DAP-seq technique for detecting DNA binding events of interacting TFs which we refer to as double DAP-seq (dDAP-seq). As illustrated in Fig. 1a, instead of expressing only one TF fused to a HaloTag as in the published DAP-seq protocol, in dDAP-seq we simultaneously expressed two TFs fused to different affinity tags, one to a Streptavidin-Binding Peptide Tag (SBPTag-TF1) and the other to a HaloTag (HaloTag-TF2). After incubating SBPTag-TF1 and HaloTag-TF2 with fragmented genomic DNA, we used HaloTag-ligand coupled magnetic beads to specifically capture potential heterodimeric complexes of HaloTag-TF2 and SBPTag-TF1, and any DNA fragments that they specifically bound. Sequencing of recovered DNA fragments revealed the locations of the heterodimer binding events throughout the whole genome. We note that dDAP-seq is particularly useful for testing heterodimerization when one or both of the TFs do not bind DNA on their own. For example, to test for heterodimerization among bZIP C/S1 family members, HaloTag-TF2 corresponded to a Group C bZIP, which could not bind DNA by itself. SBPTag-TF1 corresponded to a Group S1 bZIP, which could bind DNA as a homodimer but was not captured during the HaloTag affinity purification step.

**Fig. 1:**
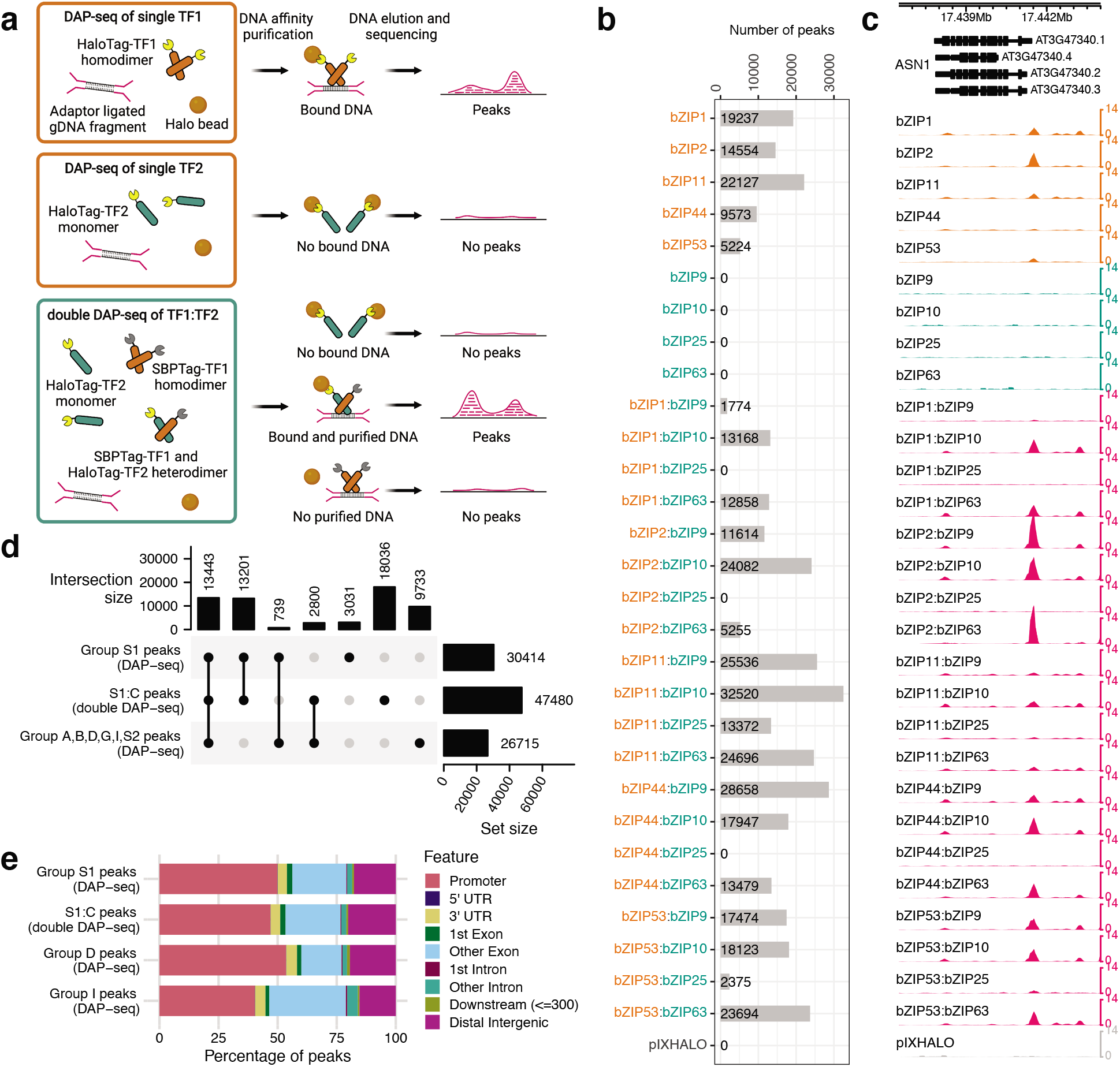
Systemic identification of bZIP C/S1 bZIP homodimer and heterodimer binding sites by DAP-seq and dDAP-seq. **a** Schematic of DAP- and dDAP-seq. Top panel: In DAP-seq, an *in vitro* expressed, HaloTag-fused TF (HaloTag-TF1) forms homodimers that bind to genomic DNA (gDNA) fragments ligated to Illumina-compatible sequencing adapters. The TF-DNA complex is purified by HaloTag ligand-coupled magnetic beads, from which the bound gDNA is eluted and sequenced. Mapping the sequencing reads to the reference genome allows identification of TF binding location as peak regions of significant read enrichment. Middle panel: Performing DAP-seq for a TF that cannot bind DNA by itself does not produce any peaks. Bottom panel: In double DAP-seq, two TFs fused to SBPTag and HaloTag separately, SBPTag-TF1 and HaloTag-TF2, are co-expressed *in vitro* and allowed to form heterodimers. HaloTag ligand-coupled magnetic beads are used to purify the complex of SBPTag-TF1:HaloTag-TF2 with the bound DNA. Although the beads can pull down the HaloTag-TF2 monomer, no peaks will be detected for them without the bound DNA. **b** Number of peaks from all pairs of C/S1 bZIPs detected by DAP-seq and dDAP-seq. Group S1 bZIPs (bZIP1, 2, 11, 44 and 53) are indicated by orange color, and Group C bZIPs (bZIP9, 10, 25 and 63) are indicated by teal color. **c** DAP-seq and dDAP-seq binding profiles of C/S1 homodimers and heterodimers at the known target gene *ASN1* (AT3G47340). **d** Upset plot comparing the peak overlap between S1 homodimers, S1:C heterodimers and bZIPs from Group A, B, D, G, I and S2. The dot plot lists the possible combinations between the different bZIP groups. The vertical bar plot at the top reports the number of peaks bound by each combination, and the horizontal bar plot on the right shows the total number of peaks bound by each bZIP group. **e** Distribution of binding sites relative to genomic features for the bZIP homodimers and heterodimers. Promoter regions were defined as ±1 kb from the TSS.

Using dDAP-seq, we comprehensively profiled the genome-wide binding events of twenty heterodimer pairs between five Group S1 and four Group C members. (Henceforth we use “C/S1” to refer to dimers among these nine bZIPs, including both homo- and heterodimers, and “S1:C” to exclusively refer to the heterodimers.) In contrast to the DAP-seq experiments for Group C bZIPs alone that found no significant binding sites, dDAP-seq of the S1:C heterodimers revealed between 1,700 to 33,000 binding sites (Fig. 1b). In the few cases where the interaction was weak or undetected, such as the interactions between bZIP25 and four S1 bZIPs (bZIP1, bZIP2, bZIP11 and bZIP44), dDAP-seq detected no stable binding signal (Fig. 1b, Supplementary Fig. 1f-i) and therefore these pairs were not included in downstream analysis. Fig. 1c shows an example of peaks produced by homodimeric DAP-seq and heterodimeric dDAP-seq binding in the promoter of *ASN1*, a known target of bZIP53. Data from dDAP-seq were highly reproducible, with Pearson correlations between replicates ranging from 0.74 to 0.99 (Supplementary Fig. 2). Complete genome-wide binding site maps are provided at http://hlab.bio.nyu.edu/?data=projects/bzip_code.

Overall, a substantial fraction of S1:C heterodimer binding sites were unique compared to those of homodimers of S1 and other bZIP groups (Fig. 1d). We compared our binding site data for fifteen S1:C heterodimer pairs (dDAP-seq) and five S1 homodimers (DAP-seq) to published, single TF DAP-seq data for 21 *Arabidopsis* bZIPs in Group A, B, D, G, I and S2^40^. Combining all these genome-wide binding site maps resulted in a total of 60,983 binding events. Among these, 13,443 peaks (22.0%) were common between the S1:C heterodimers and homodimers of other bZIP groups including S1, reflecting the conserved DNA binding preferences of the bZIP family. Among the 47,480 peaks bound by the S1:C heterodimers, 26,644 peaks (56.1%) were shared with the S1 homodimers, which may represent overlapping DNA recognition sites and redundant functions between heterodimers and homodimers. Another 2,800 (5.9%) S1:C heterodimer peaks overlapped with binding sites of other bZIP groups, while 18,036 (38.0%) were new peaks not shared with Group S1 or any other bZIP groups. The formation of unique binding sites by S1:C heterodimers indicates potentially cooperative functions among C and S1 bZIPs with a different binding specificity compared to S1 alone. The binding locations of homo- and heterodimers relative to genomic features are similar to those of other bZIP groups, with about half of all binding events located in the promoter regions (Fig. 1e).

### Genome-wide binding profiles of Group C/S1 bZIP dimers

To determine the potential tissues and developmental stages where C/S1 dimerization may occur, we examined the co-expression patterns of S1 and C genes. We observed that each pair is co-expressed in multiple tissues at different developmental stages (Fig. 2a), indicating the different S1:C heterodimers could be formed in multiple tissues. This raises the possibility that the functions of different pairs of S1:C heterodimers are redundant and suggests that expression patterns alone provide limited information about functional differences between the pairs. Thus, we could not use expression patterns alone to implicate a functional role for S1:C heterodimers in tissues.

**Fig. 2:**
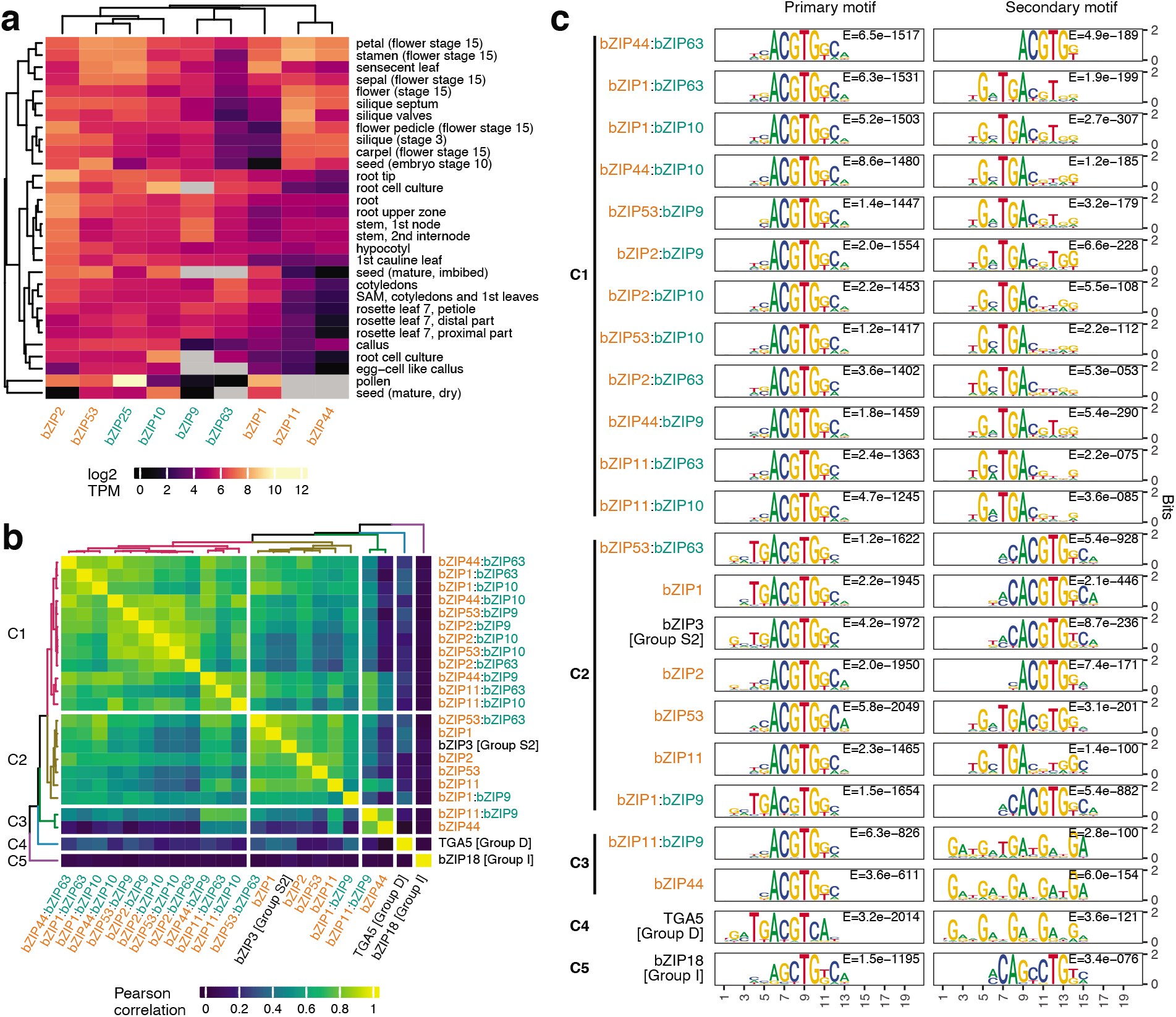
Overview of genome-wide binding correlation and DNA sequence motifs of C/S1 bZIP dimers. **a** Group S1 (orange color) and Group C (teal color) bZIPs are co-expressed in multiple *Arabidopsis* tissues at different developmental stages. **b** Hierarchical clustering using the Pearson correlation matrix of genome-wide binding profiles revealed three major clusters for C/S1 dimers (Cluster C1, C2, C3) that were separated from representative bZIPs from other groups, TGA5 (Group D) in Cluster C4 and bZIP18 (Group I) in Cluster C5. **c** PWM models of the most enriched (primary) and the second most enriched (secondary) motifs discovered from the 1,000 strongest bound peaks for each C/S1 bZIP homodimer and heterodimer. Cluster memberships correspond to those from (b).

Alternatively, we took a DNA-centric approach to narrow the context in which S1:C heterodimers may act. Since heterodimerization is known to alter DNA binding specificities for many bZIP proteins, we first investigated the relationships of DNA binding site locations among the C/S1 homodimer and heterodimers. To do this we calculated the Pearson correlation of genome-wide binding profiles between each pair of C/S1 dimers with representatives from other groups as outgroups and performed hierarchical clustering of the correlation matrix (Fig. 2b). We observed three major clusters for the C/S1 dimers that were separated from the two outgroup bZIPs, TGA5 (Group D) and bZIP18 (Group I). Cluster C1 contained twelve pairs of heterodimers, while the rest of the heterodimers and five homodimers were grouped together in Cluster C2 and Cluster C3. When we performed *de novo* motif discovery with the most enriched 1,000 peaks bound by each homo- and heterodimer, we found the most significant motifs from all the experiments shared a core ACGTG binding sequence (Fig. 2c). Peak sequences for many of the S1:C heterodimers were also enriched for a secondary motif containing the sequence TGAC, which was absent in most of S1 homodimers (Fig. 2c). For example, 397 out of the top 1,000 peaks (39.7%) for the heterodimer bZIP53:bZIP10 contained the TGAC motif. Remarkably, the patterns of occurrence of secondary motifs largely coincided with the clusters formed by genome-wide binding correlation (Fig. 2b). All the heterodimers in Cluster C1, except for bZIP44:bZIP63, had enriched TGAC secondary motifs. The homodimers and heterodimers in Cluster C2, except for bZIP11, had enriched ACGTG secondary motifs. Cluster C3, consisting of only the bZIP11:bZIP9 heterodimer and the bZIP44 homodimer, had G-rich secondary motifs. Therefore, both binding profile comparison and motif discovery results suggest that heterodimerization with Group C changes the DNA binding preference of Group S1 bZIPs, likely expanding their regulatory repertoire.

### Target genes near dDAP-seq peaks of C/S1 dimers overlap with their known targets

To determine whether the S1:C heterodimer binding events identified by dDAP-seq could regulate gene expression, we compared the differentially expressed genes (DEGs) from RNA-seq of *bzipS1* quintuple mutant rosette leaves^32^ to three sets of target genes based on DAP- or dDAP-seq binding: the S1:C heterodimer targets from dDAP-seq, S1 homodimer targets from DAP-seq, and as a control the targets of Group D and I bZIPs from DAP-seq. For each set of DAP-seq or dDAP-seq binding peaks, we calculated the target scores for all the protein coding genes in the genome using the ClosestGene method in TFTargetCaller. This method computed a score for each gene based on the overall distribution of distances between peaks to genes in a particular peak dataset and was shown to perform well in comparison to several commonly used peak-to-gene assignment methods^46^. The 322 DEGs in the *bzipS1* mutant were grouped into six clusters based on the target scores (Fig. 3a Cluster 1 to 6; Supplementary Table 1). Cluster 1 contained 27 genes that were shared targets of S1 homodimers and S1:C heterodimers but were not enriched in targets of Group D or I. Notably, 36 genes in Cluster 2 were uniquely enriched for S1:C heterodimer targets but not the targets of other groups. Cluster 3 and Cluster 4 genes displayed specific binding by Group I and D, respectively. While Cluster 5 showed weak binding for both S1 and S1:C, Cluster 6 genes showed no enrichment from any of these bZIP groups, possibly representing indirect targets. We further noted that the genes in Cluster 1-5 were mostly down regulated in the *bzipS1* mutant (Fig. 3a), suggesting that the S1 bZIPs mostly activate gene expression. To quantify the significance of association between the DAP- and dDAP-seq target scores and the up- or down-regulated gene sets in the *bzipS1* mutant, we performed a Gene Set Enrichment Analysis (GSEA)-style calculation for the target gene score and the DEGs, using the minimal hypergeometric (mHG) test that is more powerful than the standard one-sided Kolmogorov-Smirnov test in GSEA^47^. We found that the targets identified for S1 homodimers and S1:C heterodimers were the most enriched for the down-regulated DEGs, and the targets from Group D and Group I were much less significantly enriched (Fig. 3b). These results suggest that DAP- or dDAP-seq reported targets of group C/S1 dimers are transcriptionally regulated by the S1 bZIPs *in planta*, providing evidence that the S1 homodimers and the S1:C heterodimers have overlapping and unique targets.

**Fig. 3:**
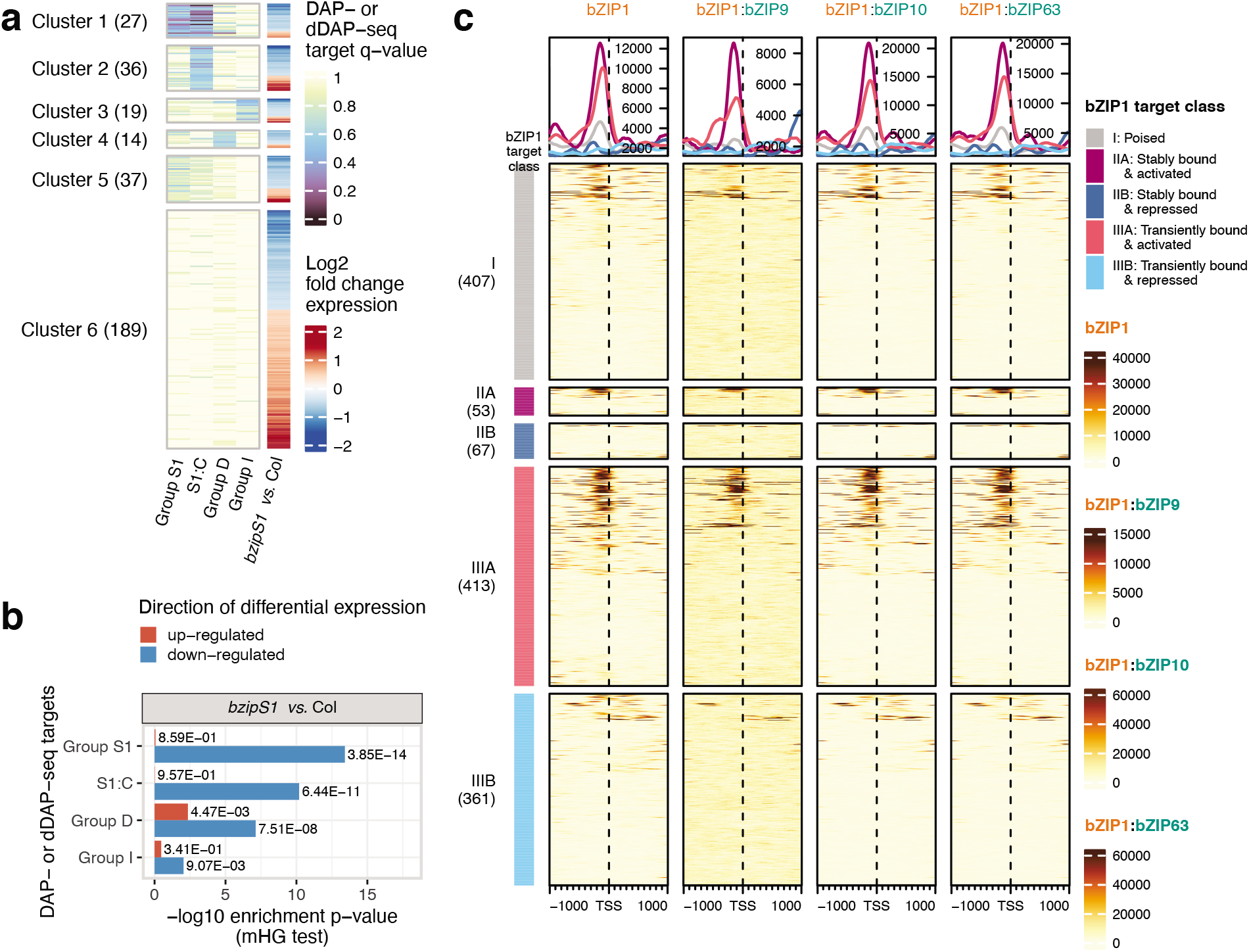
dDAP-seq identifies direct target genes of C/S1 bZIPs. **a** Distribution of DAP- and dDAP-seq target gene scores for the differentially expressed genes in *bzipS1* mutants. **b** Significance of association between DAP- and dDAP-seq target gene scores and DEGs in the *bzipS1* mutant calculated by the minimal hypergeometric (mHG) test. **c** Heatmap showing the DAP-seq and dDAP-seq binding signal of bZIP1 homodimer and heterodimers centered at the transcription start site (TSS) of bZIP1 *in vivo* targets. Left panel indicates the class of bZIP1 targets determined from ChIP-seq and time course TARGET experiments to be transiently or stably bound and activated or repressed by bZIP1.

We previously showed that single DAP-seq reported binding events were highly consistent with the *in vivo* direct binding sites identified by ChIP-seq and that target genes significantly overlapped with those obtained from TF overexpression experiments^40^. Since the S1 bZIPs can form homo- and heterodimers to bind DNA, it is unclear how the heterodimer binding found by dDAP-seq relate to the target genes found by ChIP-seq and overexpression of the S1 bZIPs. The targets for bZIP1 (Group S1), a master regulator in nitrogen response, was previously investigated by ChIP-seq and an inducible overexpression assay in protoplasts known as TARGET (transient assay reporting genome-wide effects of transcription factors)^48^. A comparison of ChIP-seq and time-course TARGET data, which implicated transient binding events, showed that bZIP1 direct target genes fell into three classes: poised (Class I), stable (Class II), and transient (Class III). Each gene in the three classes was either activated (IA, IIA or IIIA) or repressed (IB, IIB or IIIB). Using these target genes, we plotted the normalized sequencing reads from dDAP-seq within the 2 kb region centered at the transcription start sites (TSS) (Fig. 3c). For both bZIP1 homo- and heterodimers, we observed strong read enrichment in regions immediately upstream of the TSS for many of the bZIP1 targets, suggesting binding at the proximal promoter could regulate the expression of these genes. More importantly, both DAP- and dDAP-seq showed strong binding at the promoters of transient targets of Class III, which are regulated by bZIP1 overexpression but could not be detected in ChIP-seq at a single time point^48^. Furthermore, we observed stronger binding signals by both bZIP1 homo- or heterodimers at bZIP1 activated targets than at the repressed targets. The highly similar homo- and heterodimer binding profiles at the bZIP1-regulated targets suggest that those targets may also be regulated by S1:C heterodimers that contain bZIP1.

### Enriched Gene Ontology (GO) terms of the C/S1 dimer targets support their functional diversity

To compare the potential biological functions of C/S1 bZIP homodimers and heterodimers, we performed GO enrichment analysis for the top 2,000 target genes identified in DAP- and dDAP-seq based on their target scores (Fig. 4a). We observed that targets of S1 and S1:C were enriched for high level biological processes consistent with known functions of this family, such as responses to abiotic stress and light, as well as carbohydrate metabolic processes^49^. Circadian rhythm, response to water and water deprivation, and response to hormones including auxin were enriched for almost all target gene sets (Fig. 4a Cluster 1, 2 and 4), suggesting these are the conserved functions of the S1 and C groups. For a subset of the heterodimers, multiple GO terms related to hypoxia response were strongly enriched (Fig. 4a Cluster 4). Hypoxia response is important in both stress tolerance and development. For instance, internal oxygen limitation occurs in the developing seed and could regulate seed growth and physiology^50^. Our results suggest that the important role of C/S1 bZIPs in seed development^30,51^ may be mediated by hypoxia-related target genes. In fact, the functions of bZIPs in tolerance to oxidative stress can be traced back as early as algae^52^. Since hypoxia responses are also relevant in the context of plant response to submergence^53,54^, we compared the S1 and S1:C targets to genes that are differentially expressed following submergence treatment^55^ and observed a much higher level of enrichment of S1 and S1:C targets in submergence-induced genes compared to submergence-repressed genes (Fig. 4b). Interestingly, we found the targets of bZIP11 homo- and heterodimers were highly enriched for responses to auxin (Fig. 4a box labeled a), consistent with the role of bZIP11 in modulating auxin-induced transcription^56^. Taken together, the target genes identified by DAP- and dDAP-seq recapitulated important aspects of C/S1 functions.

**Fig. 4:**
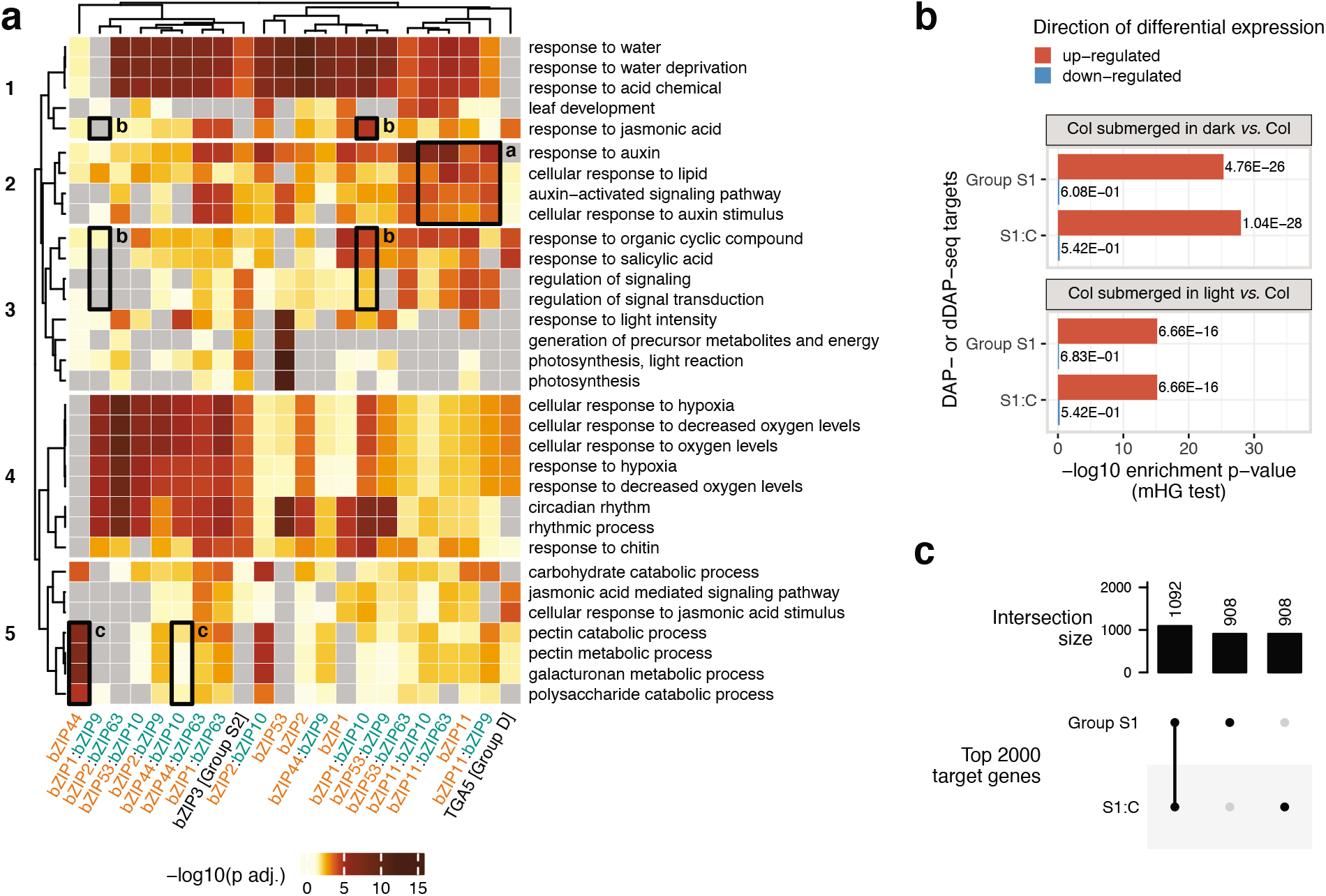
Comparison of enriched GO terms and target genes for S1 homodimers and S1:C heterodimers. **a** Enriched GO terms for top 2,000 target genes determined from DAP- and dDAP-seq peaks of C/S1 dimers and representatives from Group D (TGA5) and Group S2 (bZIP3). Grey color indicates no significant association was found for that GO term (adjusted *P* > 0.05). **b** Significance of association between DAP- and dDAP-seq targets with DEGs in response to submergence. **c** Upset plot comparing the top 2,000 target genes between the S1 homodimers and S1:C heterodimers.

Besides the common GO terms discussed above, target genes predicted for the S1 and S1:C pairs were strongly enriched in a diverse set of GO terms, showing a landscape of biological functions resulting from heterodimerization. For example, the GO terms related to responses to jasmonic acid, salicylic acid, and regulation of signaling were enriched in target genes of bZIP1:bZIP10 heterodimer but not in bZIP1:bZIP9 heterodimer (Fig. 4a boxes labeled b). bZIP44 homodimer target genes were significantly enriched in GO terms related to catabolic processes of cell wall components, which were not found for the bZIP44:bZIP10 heterodimer (Fig. 4a boxes labeled c). Overall, when we compared the top 2,000 genes targeted by the S1 homodimers and S1:C heterodimers, about half of the genes (1,092) were shared between the homodimers and heterodimers and the rest were unique (Fig. 4c). These lines of evidence support the role of heterodimerization in facilitating diverse biological functions of C/S1 bZIPs.

### Analysis of bZIP9 heterodimer targets revealed bZIP9 functions in ABA response

It is challenging to investigate the direct target genes of the Group C bZIPs as most of them do not bind DNA via homodimerization, and there is limited characterization of their biological functions due to a high degree of redundancy among members in the group. We hypothesized that dDAP-seq results for the S1:C bZIPs could help reveal the functions of Group C bZIPs and chose to explore this hypothesis by focusing on bZIP9.

bZIP9 heterodimerizes with all five S1 bZIPs^27,28^ and our dDAP-seq assay reported between 1,774 and 28,658 peaks for the five heterodimers (Fig. 1b). GO analysis of the heterodimer target genes found a diverse set of functions for the individual heterodimers, including response to water, hypoxia, and auxin stimulus (Fig. 4a). We reasoned that the functions of bZIP9 could be identified by collectively analyzing the target genes of all its heterodimers. We called peaks by merging the dDAP-seq data from the five bZIP9 heterodimer experiments and calculated the enriched GO terms for the genes near the merged peaks. The most enriched biological processes included response to abiotic stress such as water, salt and hypoxia, development, and response to hormones (Fig. 5a). One of the significantly enriched GO terms was response to abscisic acid (ABA), a plant hormone with well-characterized functions in response to water, salt, and post-embryonic development such as seed development^57–59^. We also found that the expression of bZIP9 was one of the most strongly induced by ABA among the nine C/S1 bZIPs (Supplementary Fig. 3^60,61^). Therefore, we hypothesized that bZIP9 may be involved in the regulation of ABA response, which may contribute to the responses to stress and development.

**Fig. 5:**
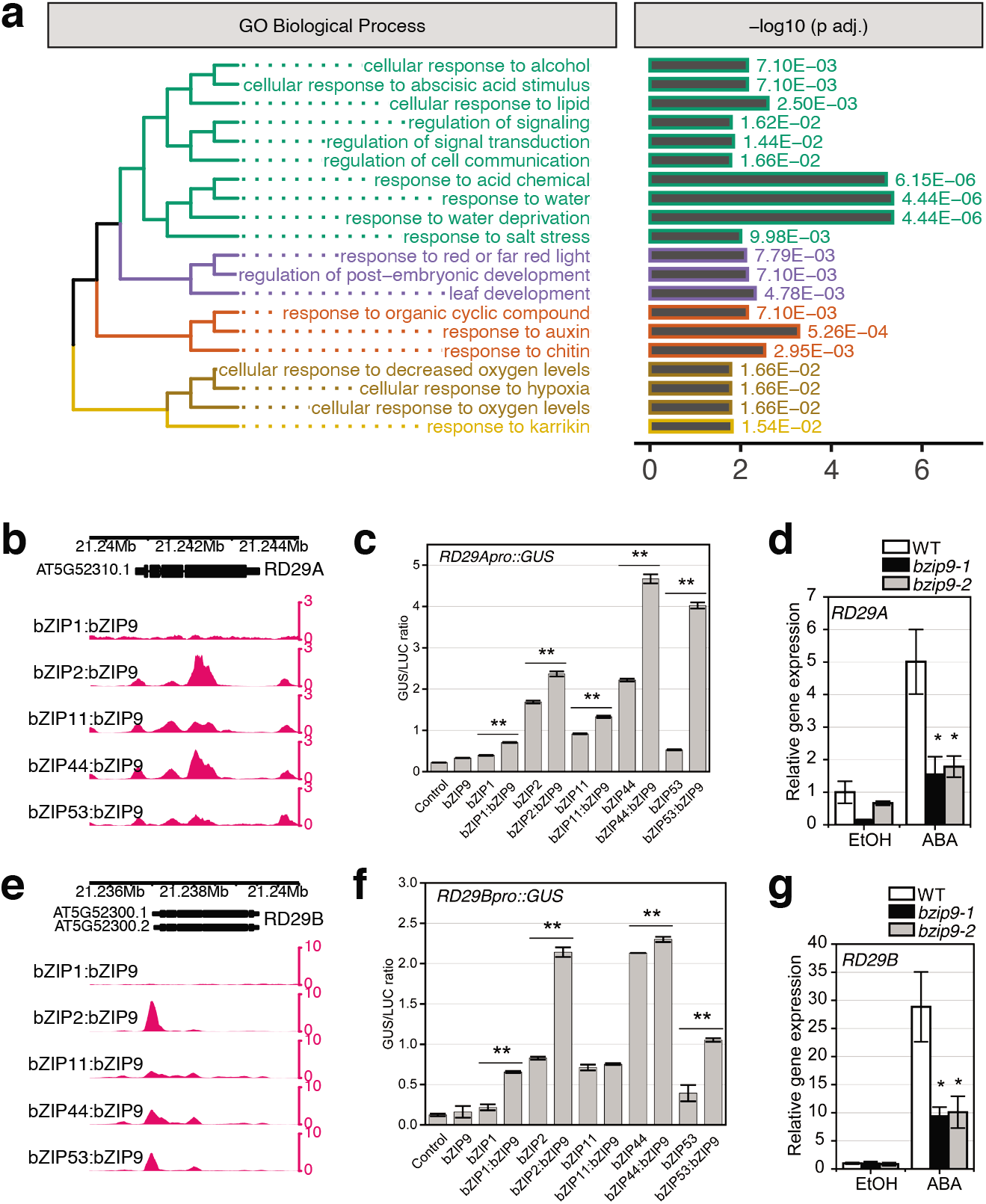
bZIP9 heterodimers have functions in ABA response. **a** Enriched GO terms for bZIPS1:bZIP9 heterodimer targets. dDAP-seq data of bZIP1:bZIP9, bZIP2:bZIP9, bZIP11:bZIP9, bZIP44:bZIP9 and bZIP53:bZIP9 heterodimers were combined to perform the GO enrichment analysis. **b, e** bZIPS1:bZIP9 heterodimer peaks located at the promoter region of ABA response marker genes *RD29A* (AT5G52300), and *RD29B* (AT5G52310). **c, f** Transient expression of bZIP9 with each bZIPS1 resulted in significantly higher expression of the *RD29A::GUS* and *RD29B::GUS* reporter compared to the bZIPS1 alone. n=3 replicates. Data are normalized to luciferase control (LUC). Bar charts represent mean ± standard error (SE). Asterisks represent statistically significant difference between bZIPS1:bZIP9 and bZIPS1 by *t*-test (*P* < 0.01). **d, g** Expression of *RD29A* and *RD29B* genes in wild type (WT) and *bzip9* mutants after treatment of 50 μM ABA for 3 h with ethanol (EtOH) as negative control. Error bars indicate SE of three independent seedling pools. Asterisks indicate statistically significant difference between *bzip9* mutants and WT in ABA-treated or control conditions by two-tailed *t*-test (*P* < 0.05).

To test our hypothesis, we started by examining bZIP9 heterodimer binding at known ABA response marker genes. *RD29A* and *RD29B* are two well-known markers of ABA response^62,63^, and we found strong binding signal at the promoters of these two genes by bZIP9 heterodimers with bZIP2, bZIP44, and bZIP53 (Fig. 5b and e). Using a transient expression assay^64,65^ where expression of a GUS reporter was driven by promoters of *RD29A* and *RD29B* (*RD29Apro::GUS* and *RD29Bpro::GUS*), we found that co-transfection of bZIP9 with individual S1 bZIPs resulted in significantly higher reporter activation than transfections by each S1 bZIP alone (Fig. 5c and 5f). Furthermore, we used qPCR to check the expression of these two marker genes in two independent T-DNA mutant lines, *bzip9-1* and *bzip9-2*, and found their expression was significantly reduced in the mutant compared to wild type following ABA treatment (Fig. 5d and g). Therefore, the DNA binding and gene expression measurements support the role of bZIP9 in the regulation of ABA response. In agreement, a Group C bZIP in wheat, TabZIP14-B, was reported to be a positive regulator of ABA and abiotic stress responses when expressed heterologously in *Arabidopsis*^66^, suggesting that the function of group C bZIP in ABA response may be evolutionarily conserved.

### bZIP53 heterodimer DNA binding specificities have sequence basis

The function of bZIP53 (Group S1) in regulating seed development has been extensively characterized^30,51^, and its heterodimerization with Group C bZIPs was shown to be an important mechanism for regulating a handful of target genes. To systematically investigate the contributions of bZIP53 heterodimers to seed development, we first identified heterodimer-specific binding sites by performing differential binding analysis comparing the bZIP53 heterodimer experiments to the bZIP53 homodimer experiment. From each pairwise comparison, we obtained between 2,143 and 9,136 peaks that were more enriched in the heterodimer compared to the homodimer (heterodimer-specific) and between 851 and 1,817 peaks that were less enriched (homodimer-specific) using adjusted *P*-value threshold of 0.05. We then compared the genes near the differentially bound peaks to those expressed in specific seed sub-regions or during specific stages in the developing seed^67^. Of the 47 sets of genes showing dominant patterns (DP) of expression in specific subregions and stages in the developing seed, the differentially bound targets for bZIP53 heterodimers were highly enriched for genes in DP9, DP12, and DP19, all three of which are specifically expressed at the mature green stage in multiple seed subregions (Fig. 6a, b; Supplementary Fig. 4a). In contrast, we did not observe enrichment for genes showing other dominant expression patterns, such as DP18 that are expressed specifically in the micropylar (Fig. 6b; Supplementary Fig. 4a). These results provide direct evidence for the role of bZIP53 heterodimers in regulating gene expression during seed maturation. Consistent with the finding that heterodimerization with bZIP10 increases the DNA binding affinity of bZIP53 at the promoter of seed maturation genes such as *2S2*^30^, we observed that the binding of bZIP53 to several seed maturation genes, such as *2S1*, *2S2* and *CRU3*, was dependent on heterodimerization with bZIP10 (Supplementary Fig. 4b and c). Therefore, bZIP53 heterodimer-specific binding events are associated with a subset of genes that have a unique spatial-temporal pattern of expression in the developing seed, illustrating the power of using dDAP-seq to investigate the biological processes regulated by S1:C heterodimers.

**Fig. 6:**
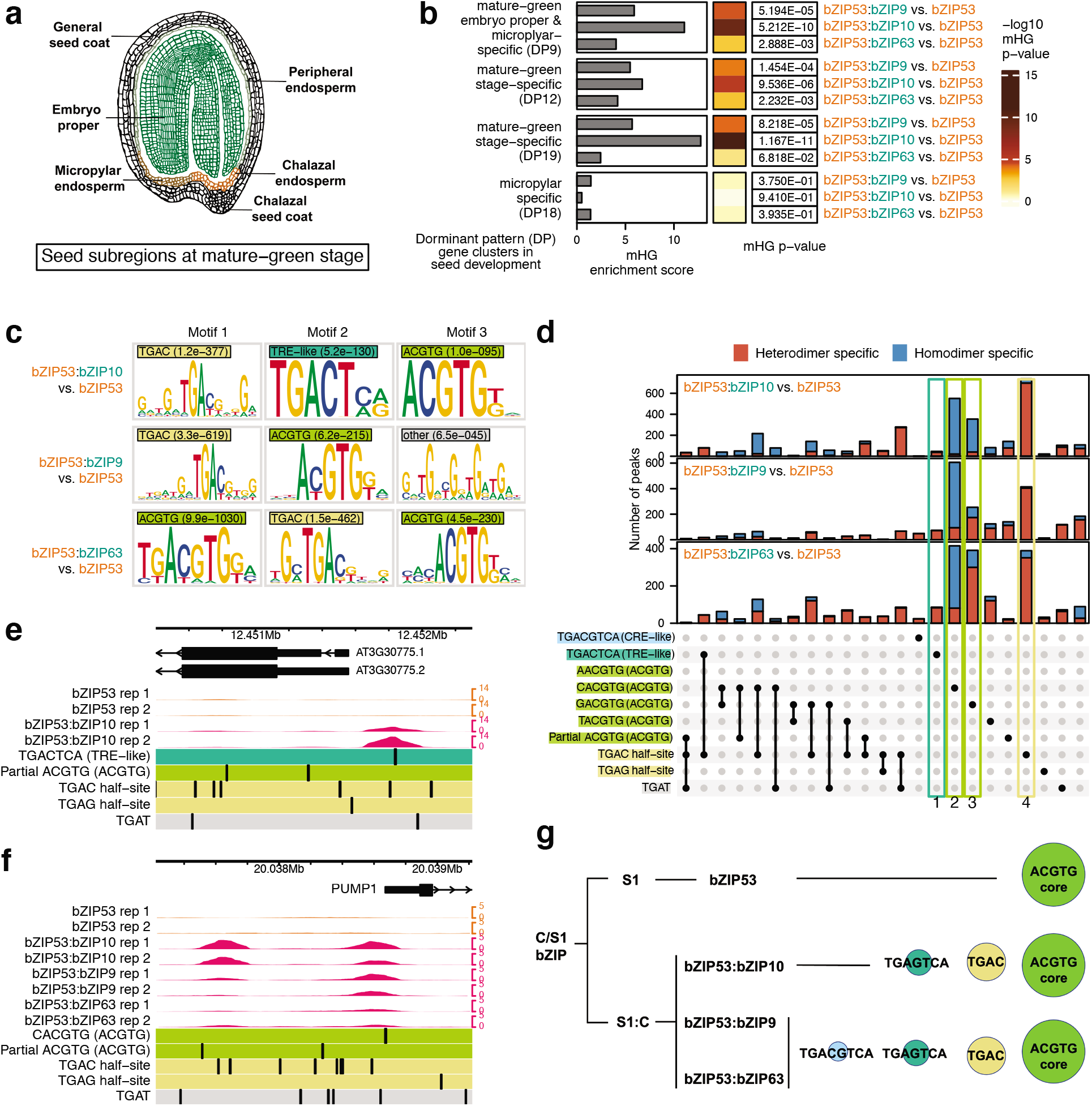
Target genes and motif differences revealed bZIP53:bZIPC heterodimer specificities. **a** Diagram of the subregions in a developing seed at the mature-green stage. **b** Significance of association determined by mHG test between the genes differentially bound by bZIP53 heterodimers and subregion-specific gene expression at the mature-green stage in the developing seed. **c** From sequences in the peaks specific to each bZIP53:bZIPC heterodimer, MEME discovered three major categories of enriched motifs: TGAC (marked by light yellow color), ACGTG (marked by pear color), and CRE-like element TGACTCA (marked by mint color). **d** KSM motif occurrences in peaks that showed increased binding comparing each bZIP53:bZIPC to bZIP53 (heterodimer-specific) or decreased binding (homodimer-specific). K-mer sequences corresponding to the motif categories in **c** are shown on the left. Each KSM motif instance in the peaks was checked for matches to the k-mer sequences, and the number of peaks containing the combinations of k-mer sequences are indicated by the bar plot at the top, separated into heterodimer- or homodimer-specific peaks. The category of sequence elements for each k-mer is included in parentheses and indicated by the background color as in **c**, and “partial ACGTG” denotes KSM matches that start or end with ACGTG. **e** A bZIP53:bZIP10 heterodimer-specific peak at the promoter region of *ProDH* (AT3G30775) contains the TRE-like element TGACTCA and a TGAC half-site. **f** bZIP53:bZIPC heterodimers have increased binding at the TGAC half-sites located near the target gene *PUMP1* (AT4G28520). **g** Model of DNA sequence specificity of bZIP53 homodimer and bZIP53:bZIPC heterodimers.

To better understand the sequence basis underlying the differential binding by homodimers and heterodimers, we sought to identify sequence motifs enriched in the homodimer- and heterodimer-specific binding events. To do this, we first extracted sequences under the peaks that are specific to each of the bZIP53:bZIPC heterodimer compared to the bZIP53 homodimer, and used these sets of sequences for *de novo* motif discovery by MEME^68^. The top three motifs enriched in the bZIP53 heterodimer-specific peaks fell into three major categories (Fig. 6c): a motif that contained a TGAC central sequence, a motif that contained a core ACGTG sequence, and a motif that contained the sequence TGACTCA. We noted that the first two motifs were similar to the motifs in Fig. 2c identified from the most enriched 1,000 peaks for each dimer, however the differential analysis uncovered additional motifs that distinguished between hetero- and homodimers. While the motif containing the core ACGTG sequence corresponded to two well-known ACGT *cis*-elements in plants, including the G-box (CACGTG) and GC-hybrid (GACGTG)^22^ sequences, the TGACTCA motif overlapped with the ACT element (ACTCAT/ATGAGT) that was previously identified for a bZIP53 heterodimer at a handful of target sites^25,29^. Binding at motifs containing a TGAC sequence that is not part of an ACGT element has not been reported for plant bZIPs. For the bZIP53:bZIP10 heterodimer, the 2,000 most heterodimer-specific peaks contained 1,601 instances of the TGAC motif and 147 instances of the TGACTCA motif, suggesting these non-ACGT elements could make a major contribution to DNA target recognition by the heterodimers. In fact, these two non-ACGT elements correspond to the high affinity binding sites previously identified for the yeast bZIP GCN4: the TRE-like element TGACTCA and the half-site TGAC. GCN4 further recognizes a third motif, the CRE-like element TGACGTCA^34–36^, which was not found among the PWM motifs enriched in bZIP53:bZIP10 heterodimer-specific binding but are strongly preferred by Group D bZIPs^40^. The presence of the relatively distinct motif categories motivated us to perform a higher-resolution, k-mer-based analysis to better capture the full complexity of the sequence specificities that may not be completely represented by PWM models used by MEME^69–71^.

From the major motif categories describe above, we determined the k-mer sequences that gave rise to each category and investigated the occurrences of these k-mer sequences in motif matches contained in the heterodimer- and homodimer specific binding events (Fig. 6d). These k-mer sequences included the CRE-like elements, TRE-like elements, the four possible ACGTG elements varying at the position before the ACGT, and the TGAC half-site with variation TGAG and TGAT that were also observed among the enriched PWM models. For finding motif matches, we used the k-mer set memory (KSM) motif representation and matching calculation^72^. Each KSM model comprise a set of aligned k-mers overrepresented in TF binding sites, capable of preserving positional dependencies and proximal flanking bases. KSM motifs were shown to more accurately model TF recognition sequences than PWM and more complex methods that incorporated high degree of positional dependencies^72^. For the three bZIP53 heterodimers, most peaks containing the TGACTCA or TGAC half-site KSM matches were heterodimer specific (Fig. 6d Box 1 and Box 4). KSM matches that contained the G-box element CACGTG were found in much higher frequency in homodimer-specific peaks (Fig. 6d Box 2). Interestingly, while bZIP53:bZIP10 was more associated with the homodimer-specific peaks that contained the GC-hybrid (GACGTG) than the heterodimer-specific peaks, the reverse was true for bZIP53:bZIP9 and bZIP53:bZIP63 (Fig. 6d Box 3). This information could not be readily obtained by inspecting the ACGTG PWMs reported by MEME (Fig. 6c), suggesting that our k-mer-based analysis was more sensitive in finding sequences targeted by homodimer- or heterodimer-specific binding. Two examples of the motif occurrences in heterodimer-specific peaks are shown in Fig. 6e and 6f. The promoter region of *ProDH* contains a known ACT-element preferred by bZIP53:bZIP10 heterodimers and we observed gain of binding signal in bZIP53:bZIP10 compared to bZIP53 (Fig. 6e). Similarly, we observed two heterodimer-specific peaks in the promoter region of *PUMP1* (Fig. 6f). The peak closest to the TSS contains a TGAC half-site and a ACGTG element near the peak summit, while the peak further upstream contains two TGAC half-sites at the peak summit but no ACGTG elements (the motif preferentially bound by bZIP53 homodimers).

Based on these results, we propose a model for the DNA binding specificities of bZIP53 homodimer and heterodimers (Fig. 6g). In general, bZIP53 homodimer mainly recognizes the classic ACGTG core motif and the TGAC half-site that is also recognized by GCN4. The bZIP53:bZIP10 heterodimer binds two types of GCN4-like motifs, the TGAC half-site and the TRE-like element TGACTCA. In addition to these motifs, bZIP53:bZIP9 and bZIP53:bZIP63 recognize a CRE-like element TGACGTCA just like GCN4. These sequence specificities provide a molecular basis for the overlapping and expanded target genes of bZIP53 heterodimers that could potentially be a general mechanism mediating the dynamic functions of C/S1 bZIPs.

### Distinct patterns of altered DNA binding specificities by C/S1 dimerization

To investigate whether the sequence patterns identified for bZIP53 heterodimer-specific binding could be generalized to other S1 bZIPs, we calculated the relative enrichment of these patterns in heterodimer-specific *vs*. homodimer-specific peaks obtained by comparing the S1:C dDAP-seq data to the corresponding S1 DAP-seq data (Fig. 7a). We observed two types of motif enrichment patterns: Type I consists of dimers of bZIP1 and bZIP44 and Type II consists of dimers of bZIP2, bZIP11 and bZIP53. For Type I, the homodimers prefer the GCN4-related motifs including the TGAC half-site, the TRE-like element TGACTCA and the CRE-like element TGACGTCA, while heterodimers prefer the ACGTG elements. Interestingly, Type II motif enrichment shows the opposite trend, where the heterodimers prefer the GCN4-related motifs and homodimers recognize the classic ACGTG elements. Therefore, although the ACGTG elements are the most enriched motifs for all the C/S1 homodimers and heterodimers (Fig. 2c) and the genome-wide binding profiles are highly similar between all the dimers (Fig. 2b), there is heterodimer-specific binding that could be explained by sequence elements that correspond to the set of sequences recognized by the yeast bZIP GCN4.

**Fig. 7:**
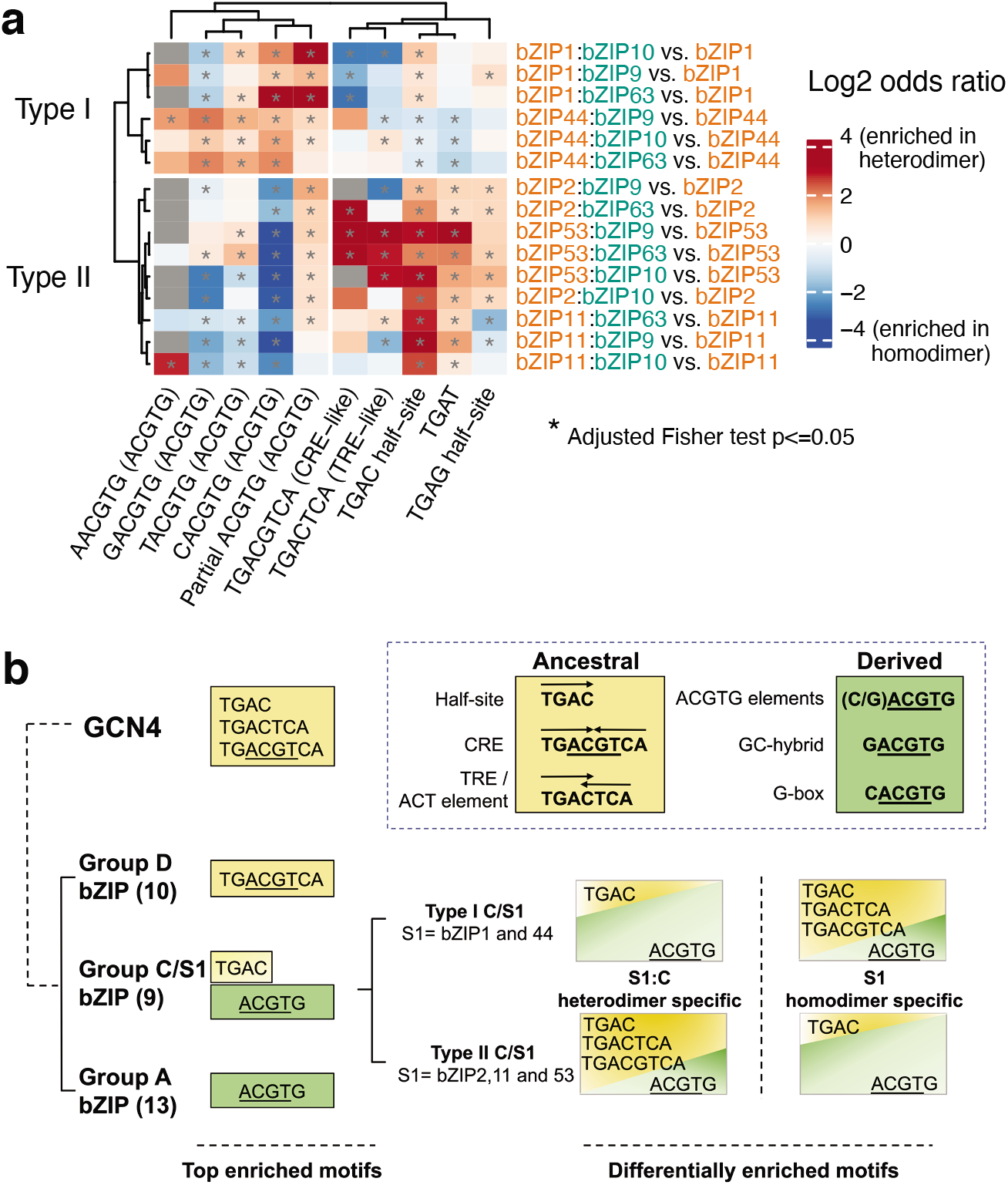
Distinct patterns of DNA binding specificities of bZIP C/S1 dimers. **a** Relative motif enrichment of differentially bound peaks for S1 homodimers and S1:C heterodimers. **b** Model for the evolution of binding specificity of the bZIP family. Plant bZIPs recognize sequences containing the core ACGTG sequence and multiple motifs targeted by the yeast bZIP GCN4. Dimers of C/S1 bZIPs have two types of sequence specificities depending on the relative enrichment of the GCN4-related motifs and the ACGTG elements. The GCN4-related motifs could represent the ancestral recognition sequences from which the ACGTG elements were derived.

## Discussion

TF interactions are key to combinatorial regulation of gene expression, so it is important to understand how interacting TFs achieve functional diversities and specificities. In this study, we upgraded the DAP-seq methodology suite to incorporate double DAP-seq, which allows mapping of genome-wide binding sites of interacting TFs in an endogenous genomic context. By expressing one or two TFs *in vitro* and comparing their bound sequences on genomic DNA, we could directly identify how TF interactions modulate binding specificities, potential target genes in the genome and biological functions. dDAP-seq is straightforward to carry out, easy to scale up, and identifies highly accurate motifs and binding events in the whole genome. Previous studies that used synthetic oligonucleotides to study binding by heterodimers across TF families and major groups of the bZIP family in human uncovered rules governing heterodimer binding in terms of changes in motif sequence and spacing constraints^10,11,73^. For the two closely related bZIP groups C and S1, the primary motifs targeted by the homo- and heterodimers are highly similar (Fig. 2c). However, since the DNA library in dDAP-seq contained the full sequence complexity and representation of the genome, we were able to uncover the differences in sequence preference between homo- and heterodimers. This system can be used to interrogate TF pairs or higher order complexes where interaction is required for DNA binding. For instance, while Group C bZIPs did not form stable homodimers in our experimental conditions, some members can form intra-group heterodimers such as bZIP10:bZIP63 and bZIP9:bZIP10^27^, which could be tested in dDAP-seq. On the other hand, if two TFs can bind DNA individually as well as when interacting with each other, such as in S1 heterodimerization, the dDAP-seq assay could be easily adjusted to allow affinity purification using the two tags sequentially ^74^. In this study, we demonstrated the sensitivity of dDAP-seq to delineate the overlapping and specific functions between the closely related C/S1 bZIPs, and we expect our approach could be applied to decipher how interacting TFs within and between families achieve the diversity and specificity of combinatorial regulation.

Evolutionary adaptation to various environmental stimuli has generally coincided with an increase in gene copy number and complexity of gene regulation^75^. Specifically, for the bZIP family that has undergone parallel expansion in plant and animal lineages, dimerization between family members is a critical mechanism to expand the DNA targeting repertoire to enable regulatory complexity and functional diversification^11,76^. Group C bZIPs in plants expanded from a Proto-C bZIP present early in the green algae Chlorophyta^77,78^. Group S1 likely originated from Proto-C or Group C by gene duplication, possessing similar protein structures to form dimers with the Group C bZIPs. The C/S1 dimers are postulated to first appear in the multicellular green algae Charophyta^77,78^. C/S1 dimerization is therefore a deeply conserved mechanism and is expected to regulate conserved target genes and biological processes. Our GO enrichment analysis of target genes for C/S1 dimers revealed biological processes correlated with two major evolutionary events in the history of plant evolution^79^. First, the processes related to response to biotic and abiotic stress, circadian rhythm, phytohormone signaling are ancient functions underlying the development of multicellularity. A second set of GO terms, which are related to leaf development and response to water and salt, are functions required for land colonization. Subsequent evolution of seed plants involved major expansion of seed genes^79^. The regulation of seed-specific gene expression could be a more recent function carried out by the heterodimers. Beyond Group C/S1, the major sub-clades of bZIPs have a different set of specific functions, for example, Group D functions in biotic response and Group I in development^26,42^. We expect dDAP-seq could be readily applied to study the DNA binding specificity of heterodimers involving these bZIPs.

Putting the two types of motif enrichment patterns we found for the C/S1 dimers (Fig. 7a) in the context of DNA sequence specificities of GCN4 in yeast and other bZIP groups in *Arabidopsis*, we arrive at an attractive hypothesis for the evolution of binding specificity of C/S1 dimers (Fig. 7b). The three motifs recognized by GCN4, the CRE-like element TGACGTCA, the TRE-like element TGACTCA and the TGAC half-site^35,80^, may represent the targets of an ancestral bZIP family in the unicellular common ancestor of fungi, plant and metazoan. The ACGTG elements are similar to the CRE-like element and were derived from this common ancestor^16^. In *Arabidopsis*, Group D bZIP recognizes the CRE-like sequence TGACGTGCA that corresponds to the ancestral sequence, while Group S and Group A bZIPs recognize the derived ACGTG element. The TGAC half-site we found for C/S1 dimers in this study has not been reported for any *Arabidopsis* bZIPs, although it is part of the consensus binding site GGATGAC identified using an *in vitro* expressed Group C bZIP from rice^81,82^, suggesting this recognition site could be conserved in higher plants. The ability of C/S1 dimers to recognize both the TGAC half-site and the ACGTG elements may mediate their distinct biological functions compared to the other bZIP groups. Furthermore, within the C/S1 Group, homodimer and heterodimer binding are distinguished by their relative preference for the ancestral and derived elements, reflecting the increased DNA binding flexibility conferred by dimerization.

The DNA binding domains of the bZIP TFs are highly conserved between plants and animals^83^, and many important insights of protein-DNA interaction have been obtained by studying this family. For plant bZIPs, the sequences flanking the ACGT elements have been extensively investigated for their role in determining DNA binding activity^22,84^. Recent analyses of published DAP-seq data, focused on the core ACGT motif, showed the flanking sequences were insufficient to determine binding^84^ and DNA shape surrounding the core motif contributed to differential binding by family members^85^. For human bZIPs, PBM experiments found that DNA binding specificities of the heterodimers were grouped into three major classes in relation to the homodimer binding sites: juxtaposing homodimer half-sites, overlapping homodimer half-sites, and emergent sites not readily inferred from the homodimer half-sites^11^. While some heterodimers bind to multiple classes of cognate sites, most display highly complex specificity landscapes that defy simple characterization. In this study, by investigating binding patterns on endogenous DNA that has the full complexity and context of binding sites in real genomes, we found that the dimers between the closely related C/S1 subgroups were distinguished by differential preference for two sets of motifs related to the GCN4 recognition sequences and ACGTG elements. As we showed previously^40^, assaying TF binding directly on genomic DNA fragments could capture the genomic properties that influence TF binding *in vivo*. It is therefore worthwhile to investigate whether general principles of DNA binding specificity of bZIP heterodimers may emerge on real genomic DNA from the complex specificity landscape determined on synthetic oligonucleotides. This knowledge is critical for more accurate prediction of how trait- or disease-associated genetic variants could impact DNA binding and combinatorial gene regulation.

## Methods

### Plant materials, treatments and phenotypic assays

For DAP-seq and dDAP-seq DNA libraires, *Arabidopsis* reference accession Col-0 (CS70000) was grown in soil at 22 °C under long-day (16h light/8h dark) conditions for three weeks. Rosette leaves were collected and flash frozen with liquid nitrogen prior to genomic DNA isolation. For ABA treatment experiments, seeds of Col-0 (WT) and *bzip9* mutants (SAIL_569_C12 and SALK_093416) were surface sterilized by 50% bleach containing 0.05% Triton-X100 for 10 minutes and washed three times by sterilized water. After 3 days of stratification at 4 °C, seeds were grown for 7 days at 22°C in long day conditions on plates containing 1 x Linsmaier and Skoog (LS) medium, pH 5.7 (Caisson laboratories, LSP03-1LT) with 0.8% agar (Caisson laboratories, A038-500GM) and 10 mg/ml sucrose (Fisher Scientific, 57-50-1). For ABA treatment, 7-day-old seedlings were transferred into the liquid LS medium with 50 μM (+/-) Abscisic Acid (ABA; PhytoTech Labs, A102) dissolved in ethanol (EtOH; 200 Proof, Pharmco, 100135). The same amounts of ethanol in the liquid LS medium were used as control. After 3-hour treatment in ABA or EtOH, the seedlings were immediately frozen in liquid nitrogen for RNA extraction.

### RNA Extraction and cDNA Synthesis

Total RNA was extracted by using RNeasy Mini Kit (Qiagen, 74104). RNA concentrations and 260/280 nm ratios were determined with a NanoDrop 2000. Total RNA was treated with RNase-Free DNase Set (Qiagen, 79254) to remove the contaminating DNA. Two μg of RNA was used for cDNA synthesis with SuperScript™ III First-Strand Synthesis System (Invitrogen, 18080051). The diluted cDNA was used as the template for Quantitative Real-Time PCR (qRT-PCR) analysis using Roche 480 LightCycler. The primers used are listed in the Supplementary Table 2.

### Standard DAP-seq and double DAP-seq experiments

#### Genomic DNA libraries preparation

The DAP- and dDAP-seq genomic DNA (gDNA) library was prepared as a standard high throughput gDNA sequencing library for the Illumina platform. Five mg gDNA eluted in 130 μl Elution buffer (10mM Tris-Cl, pH 8.5) was fragmented to an average of 200 bp using Covaris S220 Sonicator, with protocol setup “Peak Power 175.0, Duty Factor 10.0, Cycle/Burst 200, Duration 180 s, Sample temperature 4-8°C”. The reaction was precipitated by 2x volume 100% cold ethanol, washed by 70% cold ethanol and was suspended in 34 μl Elution buffer. The fragmented gDNA was ended-repaired in 50 μl reaction using the End-It DNA End-Repair Kit (Lucigen, ER81050), incubated at room temperature for 45 min. The reaction underwent ethanol precipitation as described above and the precipitated DNA was suspended in 32 μl Elution buffer and used in a 50 μl A-tailing reaction using Klenow (3’→5’ exo-) (NEB, M0212), incubated at 37°C for 30 min. The reaction underwent ethanol precipitation as described above and the precipitated DNA was suspended in 30 μl Elution buffer before adapter ligation. A-tailed gDNA fragments and 10 μl 30 μM annealed adapters were ligated in a 50 μl ligation reaction using T4 DNA Ligase (Promega, M1804) incubated at room temperature for 3 h. The DNA was precipitated and suspended in 30 μl Elution Buffer as the gDNA libraries used in DAP-seq and dDAP-seq.

#### TF protein preparation

The *pIX-Halo-bZIP* and empty *pIX-Halo* plasmids were gifts from Joseph Ecker lab. To create the *pIX-SBP-bZIP* expression plasmids, *pEntry-bZIP-ORFs* (also gifts from Ecker lab) were recombined using LR clonase (Gateway™ LR Clonase™ II Enzyme mix, ThermoFisher, 11791020) into the *pIX-SBP* vector (gifted by Mary Galli, Gallavotti lab). Plasmid DNA was extracted using the E.Z.N.A. Plasmid DNA Mini Kit I (Omega Bio-tek, D6942-01). *pIX-Halo-bZIPs* (Halo-bZIP1, 2, 11, 44, 53, 9, 10, 25 and 63) and *pIX-SBP-bZIPs* (SBP-bZIP1, 2, 11, 44, 53, 9, 10, 25 and 63) were expressed using the TNT SP6 Coupled Reticulocyte Lysate System (Promega, L4600). Expression of Halo-bZIP and SBP-bZIP proteins were confirmed by western blotting using anti-HaloTag monoclonal antibody (Promega, G9211) and anti-SBPTag mouse anti-human antibody (MilliporeSigma, MAB10764).

#### Pulldown and western blot experiments for protein-protein interaction

*In vitro* expressed Halo-protein and SBP-protein were incubated with Halo beads in 100 μl wash buffer overnight on a rotator at 4°C. The beads were washed with 100 μl cold wash buffer (PBS+0.05% NP40) for five times to remove non-binding proteins. The protein complex was eluted by heating the beads at 98°C. The SBPTag mouse anti-human antibody (MilliporeSigma, MAB10764) was used to detect the presence of SBPTag-protein in the supernatant.

#### Standard DNA affinity purification sequencing (DAP-seq)

Fifty μl protein expression reaction using TNT SP6 Coupled Reticulocyte Lysate System (Promega, L4600) containing 1000 ng *pIX-Halo-bZIP* plasmids was incubated for 3 h at 30°C. The reaction was then mixed with 10 μl Magne HaloTag Beads (Promega, G7282) and 50 μl wash buffer (PBS+0.05% NP40) on a rotator for 1 h at room temperature. The beads with protein were then washed five times on a magnet with 100 μl wash buffer to purify HaloTag-fused protein. The protein-bound beads were incubated with 100 ng adapter-ligated gDNA library in 100 μl wash buffer for 2h. The beads were then washed 5 times with wash buffer to remove unbound ligated DNA fragments. The beads were suspended in 30 μl elution buffer, heated at 98°C for 10 min, and put on ice immediately for 5 min to denature the protein and release the bound DNA fragments. 25 μl of the supernatant was used for the PCR enrichment step.

#### Double DNA affinity purification sequencing (dDAP-seq)

One-hundred μl protein expression reaction of TNT SP6 Coupled Reticulocyte Lysate System (Promega, L4600) containing 1000 ng *pIXHALO-bZIP* plasmids and 1500 ng *pIXSBP-bZIP* plasmids were incubated for 3 h at 30°C. The reaction was mixed with 20 μl Magne HaloTag beads, 100 μl wash buffer and 100 ng adapter-ligated DNA on a rotator overnight at 4°C. Halo beads were then washed 5 times on a magnet with wash buffer to purify the HaloTag-fused protein with its interacting partner. 100 ng adapter-ligated gDNA libraries and 100 μl wash buffer were added to the reaction and incubated for 6-8 h at 4°C. The beads were suspended with 30 μl elution buffer, heated at 98°C for 10 min, and put on ice immediately for 5 min to denature the protein and release the bound DNA fragments. 25 μl of the supernatant was used to PCR enrichment step.

#### Amplification of the DAP- and dDAP-seq library and sample pooling

The PCR reactions were prepared as follows: Mix 1 μl of Phusion DNA Polymerase (New England Biolabs, M0530), 10 μl of 5x Phusion HF Buffer, 2.5 μl of 10 mM dNTPs, 1 μl of Primer A (25 μM) and 1 μl of Primer B (25 μM), 25 μl of eluted DNA, add water to 50 μl. The Primer A and Primer B sequences contain unique indexes^86^ for each sample to be pooled in one sequencing run. The eluted DNA was amplified with the following PCR conditions: 98°C for 2 min, 15-19 cycles of 98°C for 15 s, 60°C for 30 s, and 72°C for 1-2 min, final extension at 72°C for 10 min. The samples were pooled and run in 1% agarose gel. The gel was cut to purify fragments from 200 bp to 600 bp using Zymoclean Gel DNA Recovery Kit (Zymo Research, D4007). The purified DNA libraries were measured by the Qubit HS ds DNA Assay Kit (ThermoFisher, Q32854) and sequenced on the Illumina platform. Relevant primer sequences are listed in the Supplementary Table 2.

### Reporter assay in protoplasts

To generate the effector plasmids used in reporter assay, the full-length CDSs of *bZIP9, bZIP1, bZIP2, bZIP11, bZIP44* and *bZIP53* were cloned in *pUC19-35S-DC* using LR reactions. Approximately 1-kb promoter fragments of *RD29A* and *RD29B* genes containing DAP- or dDAP-seq binding sites were amplified from *Arabidopsis* wild-type Col-0 genomic DNA and subcloned into *pUC19-DC-GUS* as reporter plasmids. The combinations of effector plasmids, reporter plasmids, and reference plasmids (*pUC19-35S-LUC*) were transformed into *Arabidopsis* protoplasts as described^65^. The transient reporter assay was performed as described^64^. The MUG (4-Methylumbelliferyl β-D-glucuronide) (Sigma-Aldrich, M9130) and luciferase assay system (Promega, E1500) were used to perform GUS and LUC activity assays, respectively. Relative GUS activity was calculated via normalization to LUC activity, and the data are presented as three independent biological replicates. Relevant primer sequences are listed in the Supplementary Table 2.

### DAP-seq and dDAP-seq data processing and analysis

#### Read processing, normalization and peak calling

The DAP-seq and dDAP-seq libraries were sequenced on an Illumina NextSeq 500. Adapter sequences were trimmed from the reads in the FASTQ files by Trim Galore (version 0.6.6) and Cutadapt version 3.1 with quality cutoff of 20^87^. The trimmed the reads were mapped to the *Arabidopsis* reference genome sequence TAIR10 using Bowtie2^88^ (version 2.2.9) with default parameters. Aligned reads were filtered by mapping quality score of at least 30. Peak calling was done by the GEM peak caller^89^ (version 3.3) on the filtered mapped reads with the default read distribution, TAIR10 nuclear chromosome sequences, and parameters “--f SAM ---t 1 --k_min 5 --k_max 14 --k_seqs 2000 --k_neg_dinu_shuffle --outNP --outBED --outMEME --outJASPAR --outHOMER --print_bound_seqs --print_aligned_seqs”. Peaks were called for each replicate individually or by merging the two replicates using GEM’s multi-replicate mode, with samples from experiments of empty vector *pIXHALO* and/or *pIXSBP* as control. Combined peaks for bZIP9 were called using reads from the five heterodimer dDAP-seq experiments by the GEM multi-condition mode. Each GEM run created two set of binding event calls: GEM events that were optimized for both read enrichment and centrally located motifs, and GPS events that were optimized for read enrichment only. To create the blacklist regions that contain highly enriched but artifact signals, peak calling was done for a set of negative control samples using MAC3^90^, which could find broader regions of read enrichment compared to the point-source binding events reported by GEM. The set of control samples included 18 experiments with replicates where sequence specific DNA binding was not expected to occur: DAP-seq of *pIXHALO* empty vector by HaloTag beads, DAP-seq of *pIXSBP* empty vector by SBPTag beads, double DAP-seq of *pIXHALO* empty vector and SBPTag-fused bZIP S1 by HaloTag beads, and double DAP-seq of HaloTag-fused bZIP C and empty vector *pIXSBP* by HaloTag beads. The peaks reported by MACS3 (version 3.0) for these control samples were merged and the peak regions shared by at least 5 of these control samples were used as blacklist in downstream analysis. Merged peak sets for bZIP groups were created by the program muMerge for peak overlap or target enrichment analysis^91^.

BigWig files of normalized read signals were created using the MAPQ 30 filtered alignment BAM files by the bamCoverage program in the deepTools package^92^ (version 3.5.0) with the following parameters: “--binSize 1 --normalizeUsing RPKM --ignoreForNormalization Mt Pt”. The genome browser tracks were plotted from the read normalized bigwig files by the R package karyoploteR^93^ (version 1.16.0).

#### Genome-wide binding correlation

Using the db.count method from the R/BioConductor ChIPQC package^94^ (version 1.26.0), the merged replicate GEM peaks reported for all the DAP-seq and dDAP-seq samples were combined to create a consensus peak set on which the sequencing reads were counted for each replicate, with the following arguments: minimum mapping quality score of 30 (mapQCth=30), fragment size of 200 (fragmentSize=200), each peak must be present in at least two samples (minOverlap=2), center the peaks and expand up- and downstream from the summit by 100 bp (summits=100), normalized to full library size (score=DBA_SCORE_NORMALIZED). From the consensus peak set, the regions that overlapped with the blacklist regions were removed and the regions that overlapped with the top 3,000 most enriched peaks from each replicate were kept, resulting in a filtered consensus peak set. The normalized read counts at this filtered consensus peak set were extracted for each replicate, log2 transformed, and averaged between replicates. This created a log2 normalized read count vector for each sample. Pearson correlation was calculated between all pairs of samples to create the pairwise Pearson correlation matrix. With the ComplexHeatmap package^95^ (version 2.9.4), the Pearson correlation matrix was drawn as a heatmap with hierarchical clustering dendrogram calculated using the (1-Pearson correlation) values as distances between rows and columns and the average linkage method.

#### Motif discovery and scanning

For motif discovery of the most enriched 1,000 peak sequences, the merged replicate GPS peaks were first filtered to remove peaks that overlapped with the blacklist regions. The filtered peaks were first ranked by the q-value of peak enrichment then by the fold enrichment values reported by GPS. DNA sequences were extracted from the TAIR10 reference genome for the 1,000 highest ranked peaks. MEME-CHIP^68^ (version 5.3.0) was run on these peak sequences using the following parameters: “-meme-mod anr -meme-searchsize 0 -meme-nmotifs 5”. The top two PWM motifs reported by MEME (part of MEME-CHIP) were imported into R by the universalmotif package, aligned and extended by functions in the DiffLogos package^96^, and plotted by the ggseqlogo package^97^.

The GEM peak calling process included a motif discovery step that reported the enriched motifs found as KSM motif models. The KSM models from each DAP-seq or dDAP-seq samples were scanned for matches in the TAIR10 genome sequence using the KMAC tool that was part of the GEM package^72^.

#### Mapping peaks to target genes, GO enrichment and association with known targets

To predict the target genes using DAP- or dDAP-seq peaks, merged replicate GEM peaks for each sample or muMerge peaks for the bZIP groups were first filtered to remove peaks that overlapped with the blacklist regions. Using the “ClosestGene” method in the TFTargetCaller package^46^, we calculated the target scores and q-values for all the protein coding genes annotated in Araport11. Target q-values were used when comparing between samples or bZIP groups, while gene scores were used to rank genes for enrichment of GO terms or association with known target gene sets.

For GO enrichment analysis, we took the 2,000 genes with the highest score for each sample, and used the clusterProfiler package^98^ to identify the top 5 most enriched GO categories in the Biological Process ontology annotated in the org.At.tair.db database and calculated the enrichment P-values of the GO terms in all the samples. The enrichment P-values were corrected by the Benjamini & Hochberg method and a matrix of -log10 adjusted P-values were created with enriched GO terms on the rows and DAP-seq or dDAP-seq samples on the columns and plotted as a heatmap by the ComplexHeatmap package^95^. Clustering dendrogram was obtained by hierarchical clustering of the column-centered and scaled -log10 P-value matrix with (1 – Pearson correlation) values between rows and columns as distances and the complete linkage method.

RNA-seq datasets for *bzipS1* mutant and for submergence treatment were downloaded from NCBI SRA. Quantification was performed by kallisto^99^ (version 0.45) and differential expression was determined by comparison to the appropriate wild type or mock treatment by sleuth^100^ (version 0.30). Dominant Patterns of gene expression specific to seed subregions and stages were downloaded from the supplemental data of Belmonte *et al.*^67^. The XL-mHG test^101,102^, implemented in the mhg_test function in the R package mhg, was used calculate the significance of association between target gene scores of individual samples or bZIP groups and gene sets from RNA-seq *(bzipS1* mutant, submergence treatment) or microarrays (seed subregion- and stage-specific genes).

#### Comparison of DAP-seq/dDAP-seq to targets of bZIP1 and bZIPS1

The classes of bZIP1 targets were obtained from supplementary materials of Para *et al.*^48^. Read signal at the 2 kb region surrounding the TSS of the target genes were extracted from the bigWig files and plotted in a heatmap by the EnrichedHeatmap package^103^ (version 1.21.2).

#### Analysis of differential binding between homodimers and heterodimers

MANorm2_utils^104^ (version 1.0.0) were first used to find the number of sequencing reads of all S1 DAP-seq and S1:C dDAP-seq samples contained in the GPS peak regions that occurred in at least one sample. MAnorm2 R package^104^ (version 1.2.0) was used on this count matrix to perform normalization and differential binding analysis. For each pair of one S1:C and one S1, the replicates of each experiment were normalized followed by normalization between the experiments. A mean-variance curve fit for the pair of experiments was obtained using the local fit method and all genomic intervals. Degrees of freedom were estimated from the fitted curve using only bound intervals, and differential test was done to compare the S1:C to S1 experiments using the fitted mean-variance curve and degrees of freedom. The peaks that had adjusted P-values less than 0.05 were called differentially bound. Regions that were significantly more enriched in S1:C dDAP-seq than S1 DAP-seq were called heterodimer specific and those that were significantly less enriched were call homodimer specific. For motif discovery of bZIP53 heterodimer-specific peaks, sequences from the 2,000 differentially bound peaks that had the highest fold changes when comparing each bZIP53:bZIPC to bZIP53 were used as input for MEME-CHIP^68^. For finding k-mer matches to KSM motif instances, KSM motif matches obtained previously that overlapped with 150 bp regions centered at the mid-point of all the differentially bound regions were checked for matches to the k-mer sequences in one of the motif categories, and multiple matches to the same k-mer sequence in one peak were counted as one. The significance of enrichment of the k-mer sequences in heterodimer-specific peaks was calculated by Fisher exact test function in R, comparing the number of peaks with a k-mer sequence in heterodimer-specific peaks *vs.* the number of peaks with the same k-mer sequence in homodimer-specific peaks. The significance of enrichment of the k-mer sequence in homodimer-specific peaks was calculated by the R function fisher.test, comparing the number of peaks with a k-mer sequence in homodimer-specific peaks *vs.* the number of peaks with the same k-mer sequence in heterodimer-specific peaks. Log2 odds ratios and P-values were obtained from returned values of the fisher.test function and P-values were adjusted for multiple testing by the Benjamini and Hochberg method.

## Data availability

Raw and processed DAP-seq and dDAP-seq sequencing data have been deposited in the Gene Expression Omnibus (GEO) with accession number GSE198873.

## Acknowledgments

This work was supported by the NIH award R35GM138143 and NSF Plant Genome Research Project grant IOS-1916804 to S.C.H. We thank Mary Galli and Andrea Gallavotti for the *pIXSBP* plasmid and for comments on the manuscript. We thank Ken Birnbaum and Gloria Coruzzi for comments on the manuscript. The TF expression plasmids were a generous gift from Joseph Ecker. We thank Liang Song for suggestions on experimental design. This work was supported in part through the NYU IT High Performance Computing resources, services, and staff expertise. Sequencing was performed by the NYU Genomics Core Facility with generous subsidies from the Zegar Family Foundation. This research was in part supported by the Center for Bioenergy Innovation, a Bioenergy Research Center supported by the Office of Biological and Environmental Research in the U.S. Department of Energy Office of Science. Oak Ridge National Laboratory is managed by UT-Battelle, LLC for the Office of Science of the U.S. Department of Energy under Contract Number DE-AC05-00OR22725.

## Author contributions

S.C.H. and M.L. designed experiments, performed the experiments and analysis and prepared the manuscript; W.L. identified the plant materials and did the ABA treatment experiments; W.H. performed the co-expression analysis; T.Y., W.M. and J.C. performed the reporter assay experiments and gave comments on the manuscript.

