## Supplementary Figures 1 to 4 and Legends for "Uncovering the “ZIP code” for bZIP dimers reveals novel motifs, regulatory rules and one billion years of *cis*-element evolution"

**Supplementary figures and figure legends**

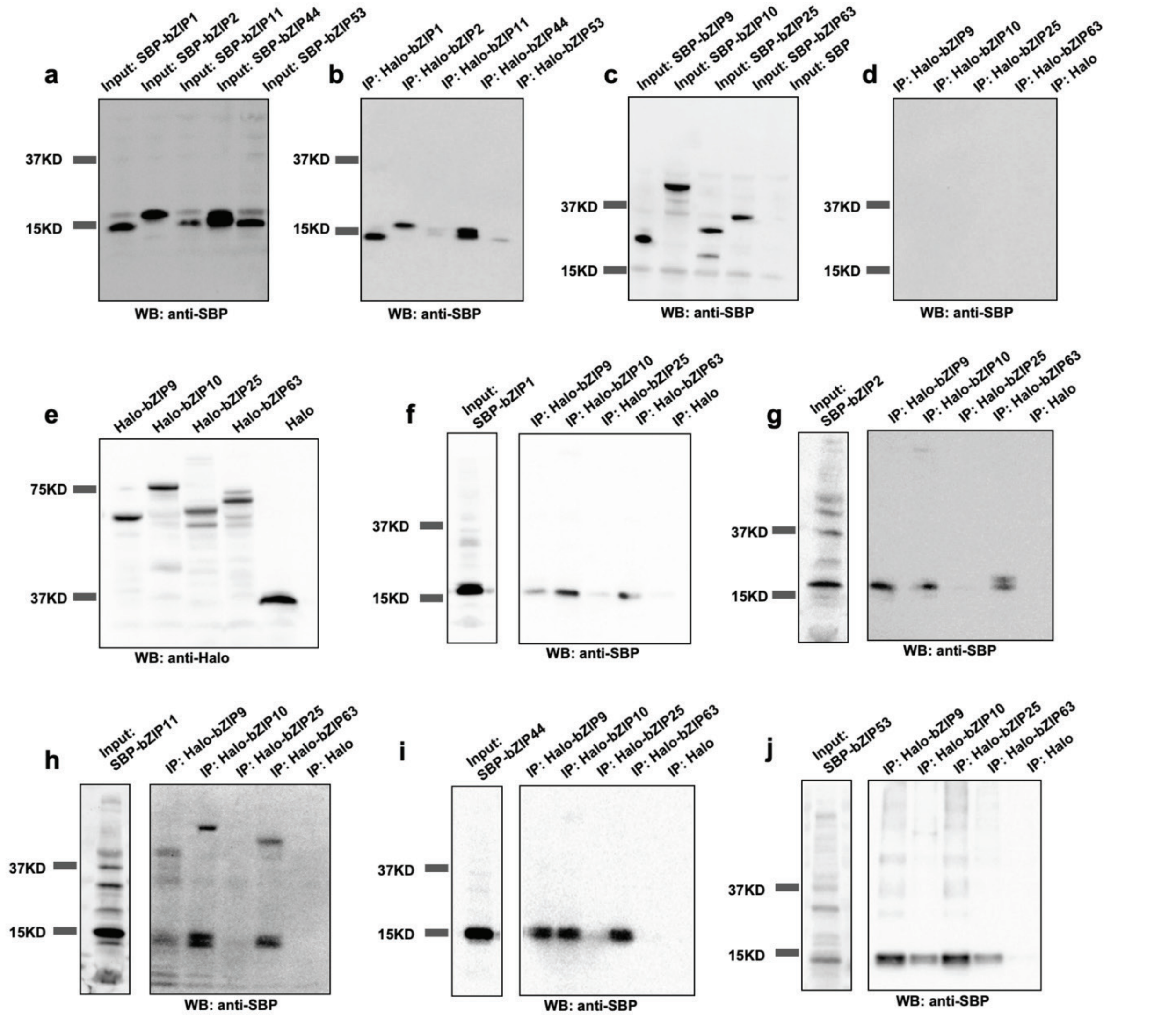


**Supplementary Fig. 1: C/S1 bZIPs form homodimers and heterodimers. a** and **b** S1 bZIP fused to SBPTag (SBP-bZIPS1) physically interact with bZIPS1 fused to HaloTag (Halo-bZIPS1) in *in vitro* pull-down assays. SBP-bZIPS1 was mixed with Halo-bZIPS1 (input samples in a) and affinity purification was carried out by HaloTag ligand-coupled magnetic beads (b). SBP-bZIPS1 was detected by an anti-SBP antibody. **c** and **d** SBP-bZIPC cannot interact with Halo-bZIPC in *in vitro* pull-down assays. SBP-bZIPC was mixed with Halo-bZIPC (input samples in c) and affinity purification was performed by HaloTag ligand-coupled magnetic beads (d). SBP-bZIPC was detected by an anti-SBP antibody. **e** Expression of Halo-bZIPC proteins and HaloTag was detected by an anti-Halo antibody. **f-j** SBP-bZIPS1 physically interacts with Halo-bZIPC *in vitro* pull-down assays. SBP-bZIPS1 was mixed with Halo-bZIPC (lanes labeled input)

and affinity purification was performed by HaloTag ligand-coupled magnetic beads (lanes labeled IP). SBP-bZIPS1 was detected by an anti-SBP antibody. The HaloTag empty vector was used as a negative control. WB: western blotting. IP: immunoprecipitation.

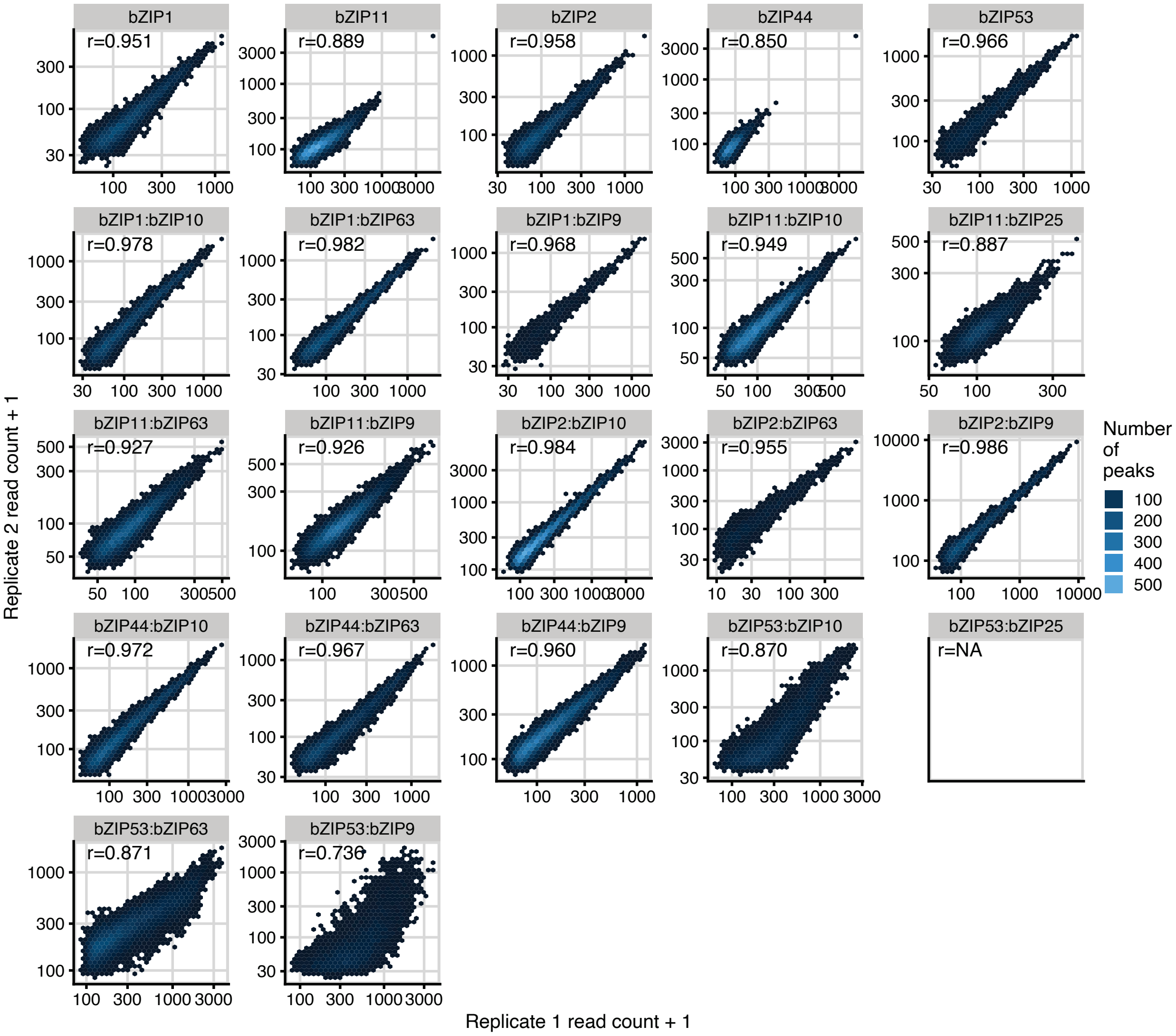


**Supplementary Fig. 2: Correlation between replicates across all the S1 DAP-seq and S1:C dDAP-seq samples.** The number of reads in peaks in replicate 1 (x-axis) and the number of reads in the same peaks in replicate 2 (y-axis) for the two replicates of each sample are plotted as hexagonal heatmaps of 2d bin counts. Pearson correlation values are indicated for each sample.

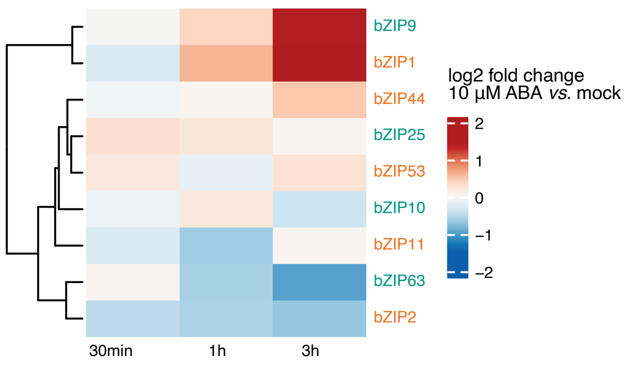


**Supplementary Fig. 3: Changes in expression of C/S1 bZIPs in an ABA treatment time course of *Arabidopsis* seedlings.** Fold changes were calculated relative to mock-treated control at the same time point.

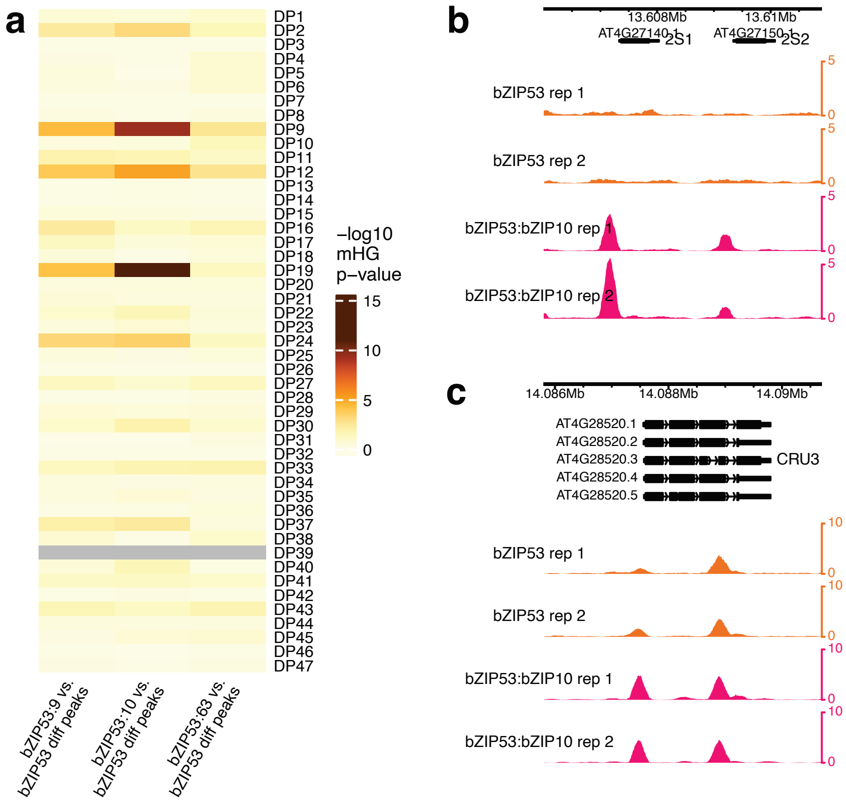


**Supplementary Fig. 4: bZIP53 heterodimer binding targets are associated with seed genes.**

**a** Minimal hypergeometric (mHG) test P-values of testing association between bZIP53 heterodimer target genes and gene sets that show distinct dominant patterns (DP) of gene expression in sub-regions at specific stages of the developing seed. **b** and **c** bZIP53:bZIP10 heterodimer binding by dDAP-seq at the promoter regions of *2S1* and *2S2* (b) and *CRU3* (c).
